# Nonparametric Interrogation of Transcriptional Regulation in Single-Cell RNA and Chromatin Accessibility Multiomic Data

**DOI:** 10.1101/2021.09.22.461437

**Authors:** Yuchao Jiang, Yuriko Harigaya, Zhaojun Zhang, Hongpan Zhang, Chongzhi Zang, Nancy R Zhang

## Abstract

Epigenetic control of gene expression is highly cell-type- and context-specific. Yet, despite its complexity, gene regulatory logic can be broken down into modular components consisting of a transcription factor (TF) activating or repressing the expression of a target gene through its binding to a *cis*-regulatory region. Recent advances in joint profiling of transcription and chromatin accessibility with single-cell resolution offer unprecedented opportunities to interrogate such regulatory logic. Here, we propose a nonparametric approach, TRIPOD, to detect and characterize three-way relationships between a TF, its target gene, and the accessibility of the TF’s binding site, using single-cell RNA and ATAC multiomic data. We apply TRIPOD to interrogate cell-type-specific regulatory logic in peripheral blood mononuclear cells and contrast our results to detections from enhancer databases, *cis*-eQTL studies, ChIP-seq experiments, and TF knockdown/knockout studies. We then apply TRIPOD to mouse embryonic brain data during neurogenesis and gliogenesis and identified known and novel putative regulatory relationships, validated by ChIP-seq and PLAC-seq. Finally, we demonstrate TRIPOD on SHARE-seq data of differentiating mouse hair follicle cells and identify lineage-specific regulation supported by histone marks for gene activation and super-enhancer annotations.

---

Context-specific regulation of gene transcription is central to cell identity and function in eukaryotes. Precision of transcriptional control is achieved through multitudes of transcription factors (TFs) that bind to the *cis*-regulatory regions of their target genes, dynamically modulating chromatin accessibility and recruiting transcription complexes in response to developmental and environmental cues^1^. Dissecting this regulatory logic is fundamental to our understanding of biological systems and our study of diseases. Over the past decades, molecular studies have elucidated the structure of TF complexes and provided mechanistic models into their function^2^. Methods based on high-throughput sequencing have enabled the genome-wide profiling of gene expression^3^, TF binding^4^, chromatin accessibility^5^, and 3D genome structure^6^. TF knockdown/knockout studies have also identified, *en masse*, their species-, tissue-, and context-specific target genes^7^. Concurrently, novel statistical approaches have allowed for more precise identification and modeling of TF binding sites^8^, and expression quantitative trait loci (eQTLs) databases now include associations that are tissue-specific^9^ and will soon be cell-type specific^10^. Yet, despite this tremendous progress, our understanding of gene regulatory logic is still rudimentary. When a TF activates or represses the expression of a gene through binding to a regulatory element in *cis* to the gene, we call such a relationship a *regulatory trio*. Despite its complexity, gene regulatory logic can be broken down into modular components consisting of such peak-TF-gene trios. In this paper, we focus on the identification of regulatory trios using multiomic experiments that jointly profile gene expression and chromatin accessibility at single-cell resolution.

Single-cell RNA sequencing (scRNA-seq) and single-cell assay of transposase-accessible chromatin sequencing (scATAC-seq), performed separately, have already generated detailed cell-type-specific profiles of gene expression and chromatin accessibility. When the two modalities are not measured in the same cells, the cells can be aligned by computational methods^11^, followed by association analyses of gene expression and peak accessibility. While these methods have been shown to align well-differentiated cell types correctly, they often fail for cell populations consisting of transient and closely similar cell states. Additionally, the alignment of cells between scRNA-seq and scATAC-seq necessarily assumes a peak-gene relationship which is usually learned from other datasets. Then, the post-alignment association analysis is plagued by logical circularity, as it is difficult to disentangle new findings from prior assumptions that underlie the initial cell alignment.

Single-cell multiomic experiments that sequence the RNA and ATAC from the same cells directly enable joint modeling of a cell’s RNA expression and chromatin state, yet methods for the analysis of such data are still in their infancy. Almost all existing methods for detecting and characterizing regulatory relationships between TF, regulatory region, and target gene rely only on marginal relationships, i.e., associations between two of the three entities without conditioning on the third. For example, Signac^12^ and Ma *et al*.^13^ use marginal associations between peaks and genes to identify putative enhancer regions, while Signac^12^ and Seurat V4^14^ link differentially expressed TFs to differentially accessible motifs across cell types. Such pairwise marginal associations are sometimes examined manually using low-dimensional embedding. One exception is PECA^15^, which uses a parametric model to characterize the joint four-way relationship between TF expression, regulatory site accessibility, chromatin remodeler expression, and target gene expression. Although PECA was designed to be applied to matched bulk transcriptomic and epigenomic data, such joint modeling concepts could potentially be very powerful for single-cell multiomic data. In this paper, we propose a nonparametric approach as an alternative to PECA’s parametric model, thus allowing for robustness and computational scalability.

As we will show through examples, context-specific gene regulation, such as cell-type-specific regulation, may be masked in marginal associations. For example, associations between a TF and its target gene may be apparent only conditional on the accessibility of its binding site. Or, associations between the accessibility of an enhancer and its target gene may be apparent only after accounting for the expression of certain transcription factors involved in, but not sufficient for, the remodeling of the enhancer region. The identification and characterization of such context-specific relationships are relevant, for example, in the interpretation of GWAS results, where marginal pairwise associations between ATAC peaks and gene expression have had limited success in linking disease-associated SNPs to genes^16^.

We explore in this paper the use of higher-order models that interrogate conditional and three-way interaction relationships for the identification of regulatory trios. First, as proof of principle, we show that a simple model that integrates TF expression with *cis*-peak accessibility significantly improves gene expression prediction, as compared to a comparable model that utilizes peak accessibility alone. We present TRIPOD, a computational framework for *t*ranscription *r*egulation *i*nterrogation through nonparametric *<u>p</u>*artial association analysis of single-cell multi*<u>o</u>*mic sequencing *<u>d</u>*ata. TRIPOD detects two types of trio relationships, which we call conditional level 1 and conditional level 2, through robust nonparametric tests that are easy to diagnose. TRIPOD’s nonparametric approach for the identification of conditional associations avoids assumptions of linearity of relationships and normality of errors, allowing for better adjustment for confounding. Thus, given a multiome experiment that measures RNA expression and chromatin accessibility for the same cells at single-cell resolution, TRIPOD outputs, for a list of transcription factors, their putative regulatory gene targets and the *cis* regions where they putatively bind to regulate each gene. This allows the prioritization of regulatory relationships for downstream analyses. We also develop a novel influence measure that allows the detection and visualization of cell states driving these regulatory relationships, applicable to data consisting of discrete cell types as well as continuous cell trajectories.

We first apply TRIPOD to single-cell multiomic data of human peripheral blood mononuclear cells (PBMCs) and compare the regulatory trios detected to relationships detected through marginal associations. We show that the detections are coherent with the vast amounts of existing knowledge from enhancer databases, bulk cell-type-specific chromatin immunoprecipitation followed by sequencing (ChIP-seq) experiments, tissue-specific TF knockdown/knockout studies, and *cis*-eQTL studies, but that conditional and marginal models identify different sets of relationships. We next apply TRIPOD to the interrogation of lineage-specific regulation in the developing mouse brain, where relationships detected by TRIPOD are compared against those derived from existing ChIP-seq and proximity ligation-assisted ChIP-seq (PLAC-seq) data. Here, TRIPOD identifies known trio relationships, as well as putative novel regulatory crosstalk between neuronal TFs and glial-lineage genes. We also apply TRIPOD to SHARE-seq data on mouse hair follicle cell differentiation to illustrate trio detection and influence analysis in data collected from different protocols. Through these analyses, we demonstrate how to harness single-cell multiomic technologies in the study of gene regulation and how the data from these technologies corroborate and complement existing data.

## Results

### A simple interaction model between TF expression and peak accessibility improves RNA prediction

To motivate our methods, we start with a simple prediction-based analysis, comparable to that done by existing methods^11^. We benchmarked against: (i) Signac^12^ and Cicero^17^, which predict gene expression by the gene activity matrix derived from the sum of the ATAC reads in gene bodies and promoter regions; (ii) MAESTRO^18^, which predicts gene expression using a regulatory potential model that sums ATAC reads weighted based on existing gene annotations; and (iii) sci-CAR^19^, which predicts gene expression by a regularized regression on coverage of individual peaks nearby. We compared the predictions derived from these methods to that of a regularized regression model, where for predictors, peak accessibilities are replaced by products between peak accessibilities and TF expressions. Only peaks within a certain range of the gene’s transcription start site (TSS) and only interactions between TFs and peaks containing high-scoring binding motifs for the TFs are considered. We refer to this model as the peak-TF LASSO model. Since this model is prediction-based, we do not expect the peak-TF pairs selected by LASSO to necessarily have a causal regulatory relationship to the gene. Comparing this model to (i)-(iii) allows us to assess whether the peak-TF interaction terms are informative for gene expression. To avoid overfitting, we performed out-of-fold prediction and adopted independent training and testing sets. See Methods for details.

We analyzed single-cell multiomic datasets from different human and mouse tissues generated by different platforms – PBMC by 10X Genomics, embryonic mouse brain by 10X Genomics, mouse skin by SHARE-seq^13^, and adult mouse brain by SNARE-seq^20^. Data summaries are included in Supplementary Table 1; reduced dimensions via uniform manifold approximation and projection (UMAP)^21^ are shown in Fig. 1a and Supplementary Fig. 1, 2a. To mitigate the undesirable consequences of sparsity and stochasticity in the single-cell data, we clustered cells to form metacells^14^ and pooled gene expression and chromatin accessibility measurements within each metacell.

**Fig. 1.**
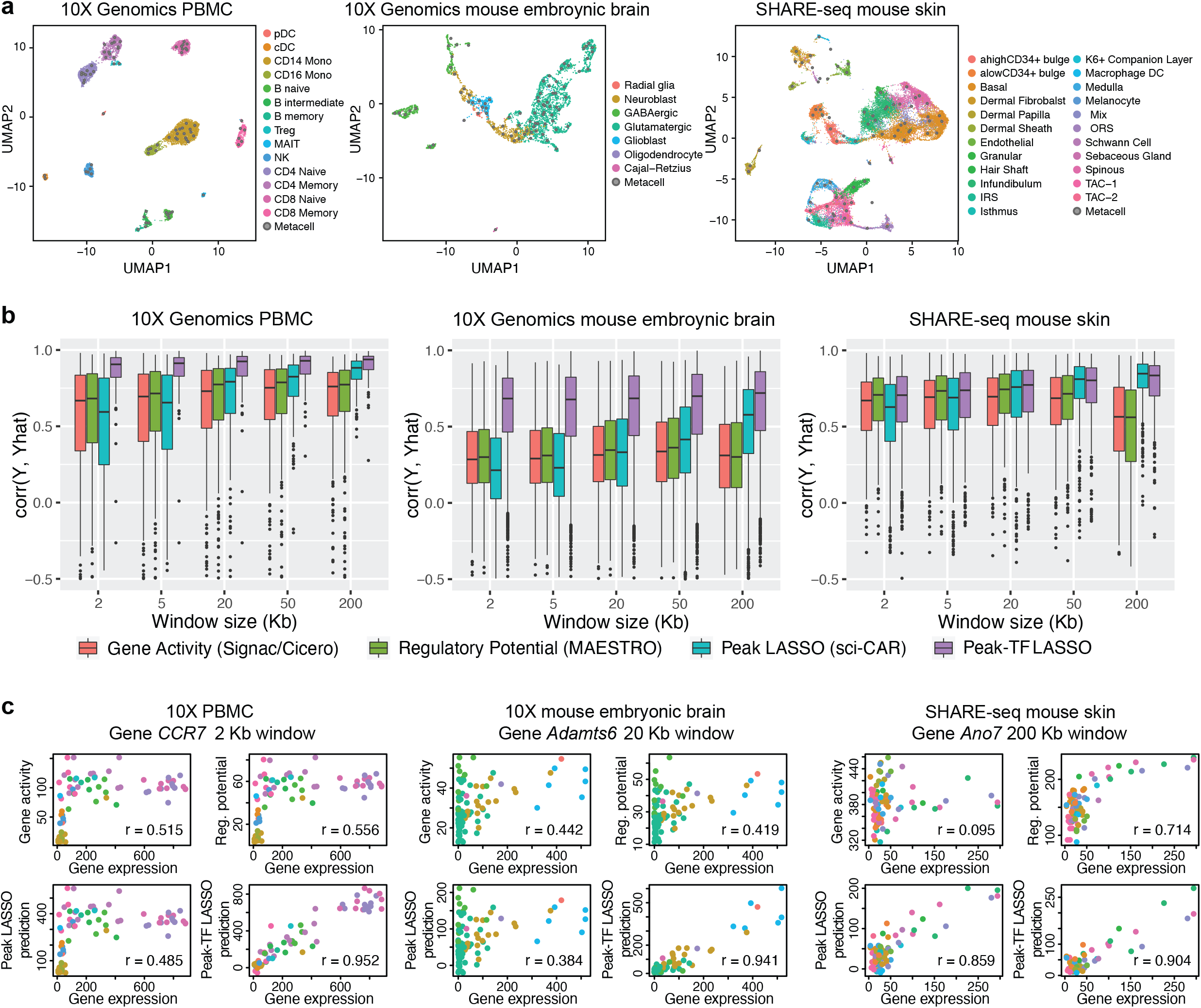
Interaction between TF expression and peak accessibility improves RNA prediction accuracy. **a**, UMAP embedding of 10x Genomics PBMC (left), 10x Genomics embryonic mouse brain (center), and SHARE-seq mouse skin (right) cells from single-cell RNA and ATAC multiomic sequencing. Cell-type labels were transferred from existing single-cell references or curated based on marker genes, motifs, and peaks; metacells were constructed to mitigate sparsity and stochasticity. **b**, Genome-wide distributions of Pearson correlations between observed and leave-one-out predicted RNA expression levels, with varying window sizes. Predictions are from gene activity, regulatory potential, peak LASSO regression, and peak-TF LASSO regression. **c**, Predicted and observed RNA expression levels for highly variable genes, *CCR7, Adamts6*, and *Ano7*, from the three datasets, respectively.

Our results show that, across window sizes, the peak-TF LASSO model significantly improves prediction accuracy across the transcriptome (Fig.1b), with examples of specific genes shown in Fig. 1c. This improvement in prediction accuracy holds true when an independent dataset is used for validation (Supplementary Fig. 3). For the SNARE-seq data^20^, sequencing depth is substantially shallower (Supplementary Fig. 4), thus the improvement of the peak-TF LASSO model is diminished but still evident (Supplementary Fig. 2b). This demonstrates that the product of TF expression and peak accessibility significantly improves RNA prediction accuracy beyond simply using peak accessibility, offering strong empirical evidence of three-way interaction relationships between TF expression, peak accessibility, and target gene expression that can be extracted from such multiomic experiments. However, we will not rely on coefficients from the LASSO model to screen for such trios, as their significance is difficult to compute due to the hazards of post-selection inference^22^. Additionally, accessibility of peaks and expression of TF affecting the same gene are often highly correlated, in which case LASSO tends to select the few with the highest associations and ignore the rest. In such cases, we believe it is more desirable to report all trios.

### TRIPOD for the detection of peak-TF-gene trio regulatory relationships by single-cell multiomic data

We propose TRIPOD, a nonparametric method that screens single-cell RNA and ATAC multiomic data for conditional associations and three-way interactions between the expression of a TF *t*, the accessibility of a peak region *p* containing the TF’s motif, and the expression of a putative target gene *g* within a pre-fixed distance of peak *p* (Fig. 2a). Existing methods^12-14^ screen for marginal associations either between the TF and the peak or between the peak and the target gene. However, three-way relationships may be complex: When a TF binds to a *cis*-regulatory region to affect the expression of a gene, it can do so in multiple ways, leading to different patterns in the data. The TF could be directly responsible for opening the chromatin of the enhancer region, facilitating the binding of other TFs that recruit the RNA polymerase. In such cases, expression of the TF is likely to be marginally correlated with the accessibility of the enhancer region, but its correlation with the expression of the target gene may be masked due to confounding of other involved TFs. Alternatively, the TF may not be directly responsible for chromatin remodeling but may bind to already accessible chromatin in recruiting other TFs or the RNA polymerase. In such cases, expression of the TF may not be highly correlated with the accessibility of the enhancer region or with the expression of the target gene. When marginal associations are masked, evidence for binding of the TF at the peak in the regulation of a gene can be inferred from partial associations: (i) with the peak open at a fixed accessibility, whether cells with higher TF expression have higher gene expression; and (ii) with the TF expression fixed at a value above a threshold, whether cells with higher peak accessibility have higher gene expression. To identify such conditional associations without making linearity assumptions on the marginal relationships, TRIPOD matches metacells by either their TF expressions or peak accessibilities (Fig. 2b): for each matched metacell pair, the variable being matched is controlled for, and differences between the pair in the other two variables are computed. Then, across pairs, the nonparametric Spearman’s test is used to assess the association between the difference in target gene expression Δ*Y*_*g*_ and difference in the unmatched variable (i.e., Δ*Y*_*t*_ if the cells were matched by *X*_*p*_, or Δ*X*_*p*_ if the cells were matched by *Y*_*t*_). We call this the “conditional level 1 test.”

**Fig. 2.**
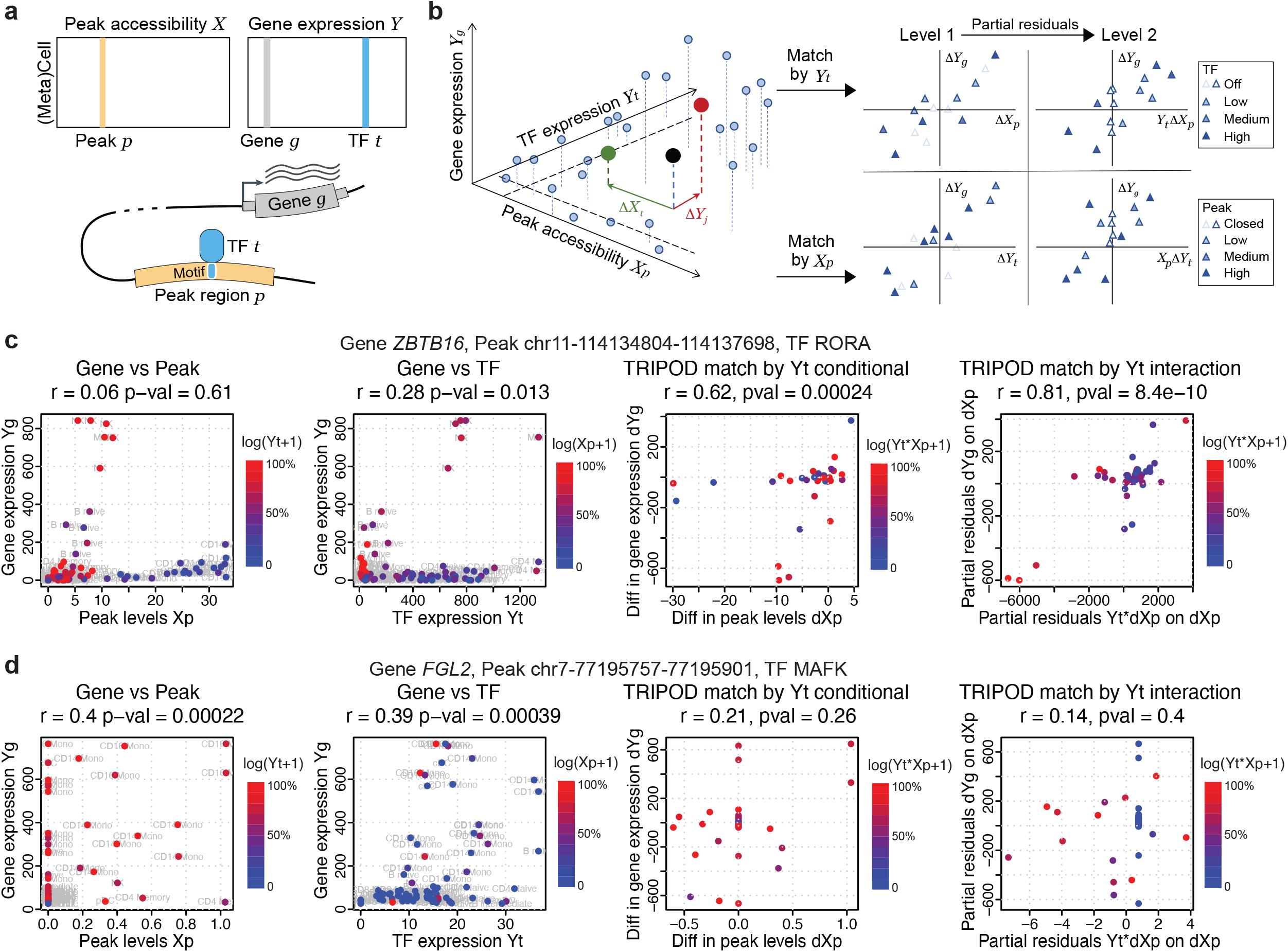
TRIPOD infers peak-TF-gene trio regulatory relationships using single-cell multiomic data. **a**, Data input and schematic on a peak-TF-gene trio. **b**, Overview of TRIPOD for inferring regulatory relationships. TRIPOD complements existing methods based on marginal associations by identifying conditional associations through matching by TF expression or peak accessibility. **c**, An example trio identified by TRIPOD, but not by the marginal associations due to the heterogeneity of cell-type-specific regulations. **d**, An example trio identified by the marginal associations, but not by TRIPOD. The peak and TF are significantly linked to the gene, yet they act through other TF and peak, and thus the regulatory trio is insignificant. The points represent metacells (left two panels) and pairs of matched metacells (right two panels). Genomic coordinates for the peaks are from hg38.

For illustration, consider the metacell denoted by the black point in Fig. 2b: If we were to match by peak accessibility, this metacell would be matched to the metacell colored in red. We would then compute Δ*Y*_*t*_, the difference between TF *t* expressions of the matched pair. If we were to match by TF expression, the black dot would be matched to the metacell in green, and we would compute Δ*X*_*p*_, the difference in peak *p* accessibility for this pair. In either case, we would compute Δ*Y*_*g*_, the difference in gene *g* expressions between the pair. We would then mask those metacell-pairs whose values, for the variable being matched, are too low (i.e., those pairs where the TF is off or the peak is closed). Then, Δ*X*_*p*_ or Δ*Y*_*t*_, together with Δ*Y*_*g*_, would be submitted for level 1 test. We call such a triplet of TF, peak, and target gene a “regulatory trio.”

Even stronger evidence for a regulatory trio could be claimed if the *degree* of association between the pairwise differences depends on the matched variable. For example, we would tend to believe that TF *t* binds to peak *p* to regulate gene *g* if, in cells with high expression of TF *t*, an increase in peak *p* accessibility yields a much larger increase in gene *g* expression, as compared to in cells with low expression of TF *t*. One could screen for such interactions by matching by either TF *t* or peak *p* accessibility. TRIPOD screens for such interaction effects through a “conditional level 2 test”, which assesses the association between Δ*Y*_*g*_ and the product of the matched variable with the difference in the unmatched variable, after taking partial residuals on the difference in the unmatched variable. In summary, TRIPOD categorizes each identified trio relationship as supported by marginal association, association between peak and gene conditioned on TF expression, and/or association between TF and gene conditioned on peak accessibility. The conditional relationships are further categorized to level 1 or level 2, with level 2 indicative of a stronger relationship exhibiting multiplicative interaction effects between TF expression and peak accessibility.

For significant trios, TRIPOD further carries out a sampling-based influence analysis, where phenotypically contiguous sets of metacells are held out to measure their influence on the estimated coefficients. The corresponding cell types/states that lead to significant deviations from the null upon their removal have high influence scores, which can be used to identify cell types/states that drive a regulatory relationship.

To highlight the differences between TRIPOD and existing methods based on marginal associations, we show two canonical examples (Supplementary Fig. 5) where the two approaches disagree. Fig. 2c outlines a significant trio detected by TRIPOD’s level 2 testing, yet the marginal peak-gene and TF-gene associations were insignificant. It turns out that a subset of cells with high peak accessibility {*X*_*p*_} have close-to-zero TF expressions {*Y*_*t*_}, and, meanwhile, another subset of cells with high TF expressions {*Y*_*t*_} have close-to-zero peak accessibilities {*X*_*p*_}. In these cells, either the peak is closed, or the TF is not expressed, and this leads to the target gene not being expressed, which masks the marginal associations. The high peak accessibility and TF expression in these cells, which act through other regulatory trios, cancel out when we consider the interaction {*X*_*p*_ × *Y*_*t*_}, leading to a significant interaction term detected by TRIPOD. Conversely, Fig. 2d outlines another trio, whose marginal associations were significant, yet TRIPOD did not detect significant conditional associations from either level 1 or level 2 testing. In this case, with almost constant TF expression, the large difference in peak accessibility leads to a small difference in target gene expression. Meanwhile, the cells that drive the significantly positive correlation between {*Y*_*g*_} and {*Y*_*t*_} have almost zero values for {*X*_*p*_}. Both observations suggest that this peak has little to do with the regulation of the target gene *FGL2* by this specific TF MAFK. Notably, we do not claim that the significantly linked peaks and TFs through marginal association are false positives, but rather this specific trio is insignificant (i.e., the peak and TF may act through other TF and peak, respectively). In summary, TRIPOD puts peak-TF-gene trios into one unified model, complementing existing methods based on marginal associations and allowing for simultaneous identification of all three factors and prioritization of a different set of regulatory relationships.

### TRIPOD identifies three-way regulatory relationships in PBMCs with orthogonal validations

We first applied TRIPOD to identify regulatory trios in the 10k PBMC dataset. Cell-type labels for this dataset were transferred from a recently released CITE-seq reference of 162,000 PBMC cells measured with 228 antibodies^14^. After quality control, we kept 7790 cells from 14 cell types pooled into 80 metacells, 103,755 peaks, 14,508 genes, and 342 TFs; the UMAP reduced dimensions are shown in Supplementary Fig. 1a. Distribution of the number of peaks 100kb/200kb upstream and downstream of the TSS per gene, as well as distribution of the number of motifs per peak, are shown in Supplementary Fig. 6.

As a proof of concept, we first illustrate two trios where the frameworks agree, identified by level 1 conditional testing (regulation of *CCR7* by LEF1; Fig. 3a) and level 2 interaction testing (regulation of *GNLY* by TBX21; Fig. 3b). From the influence analyses, TRIPOD identified B and T cells as the cell types where LEF1 regulates *CCR7*, and natural killer (NK) cells as the cell types where TBX21 regulates *GNLY*. These cell type-specific regulatory relationships are corroborated by motif’s deviation scores using chromVAR^23^ (Fig. 3) and the enrichment of Tn5 integration events in the flanking regions using DNA footprinting analyses^12^ (Supplementary Fig. 7e). Unlike chromVar and DNA footprinting analyses, which only give genome-wide average enrichments, TRIPOD significantly enhances the resolution by identifying the specific *cis*-regulatory regions that the TFs bind for the regulation of target genes.

**Fig. 3.**
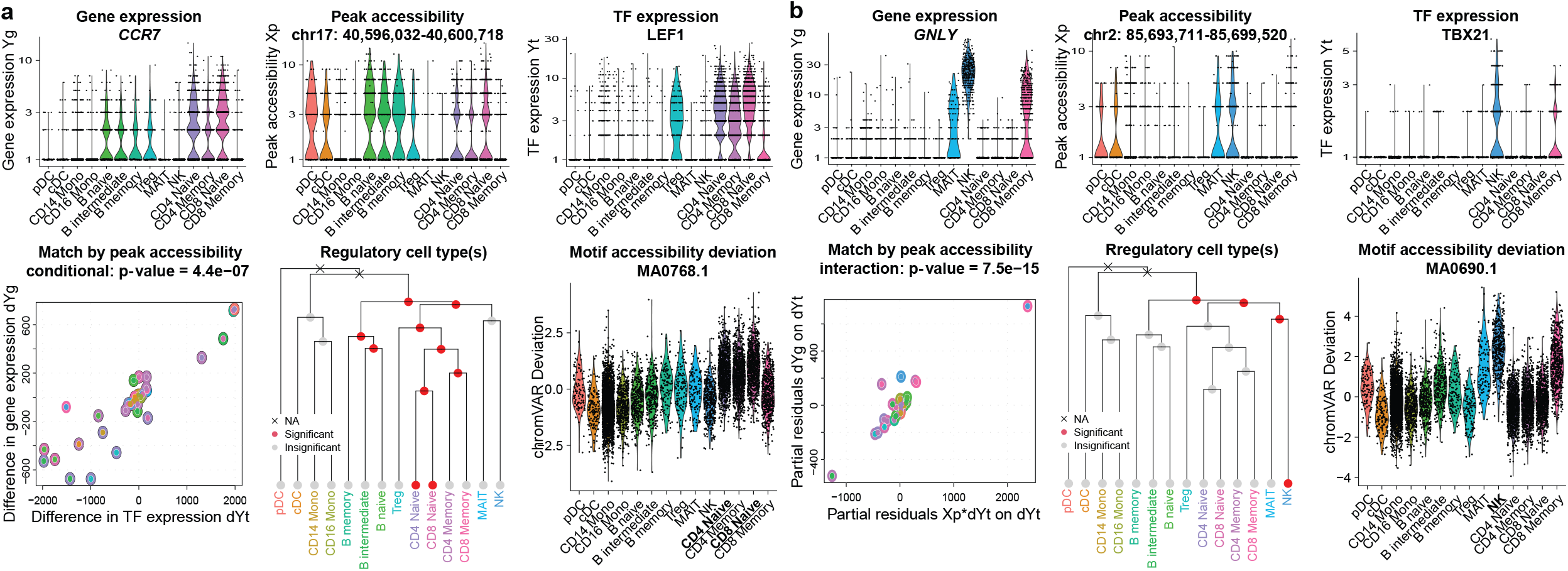
Examples of trio regulatory relationships in PBMC single-cell multiomic dataset. **a-b**, Example trios identified by TRIPOD. Violin plots show cell-type-specific distributions of gene expression, peak accessibility, and TF expression. Scatterplots show TRIPOD’s level 1 and level 2 testing, respectively. Inner and outer circles around the points are color-coded based on the cell types of the matched metacells. Hierarchical clustering is performed on RNA expression levels of highly variable genes. Red/gray circles indicate whether removal of the corresponding branches of metacells significantly changes the model fitting; crosses indicate that removal of the groups of metacells resulted in inestimable coefficients. Genomic coordinates for the peaks are from hg38.

Results from TRIPOD and marginal association tests overlap but, as expected, exhibit substantial differences (Supplementary Fig. 8). The previous section showed example trios where the two frameworks disagree. Additionally, results from TRIPOD’s matching scheme and those from random matching also overlap but exhibit substantial differences, both on the global scale (Supplementary Fig. 9) and for each gene (Supplementary Fig. 10). Notably, for the two counterexamples discussed in the previous section, random matching could not identify the masked positive trio in Fig. 2c, yet it retained significance for the negative trio shown in Fig. 2d, in a similar fashion to marginal testing (Supplementary Fig. 11). Genome-wide *p*-value distributions from TRIPOD’s two levels of testing under the null with permuted peak accessibility and TF expression are shown in Supplementary Fig. 12, indicating that TRIPOD’s framework has good type I error control. A master output of significant associations with Bonferroni correction is shown in Supplementary Table 2, with scatterplots and pairwise correlations of genome-wide *p*-values from different testing schemes shown in Supplementary Fig. 13.

To our best knowledge, no experimental technique can directly validate three-way regulatory relationships at high resolution with high throughput. Therefore, we performed validation and benchmarking by harnessing existing databases and orthogonal sequencing experiments that interrogate each pairwise relationship among the three factors (Table 1). The rationale is that true regulatory relationships should show enrichment in all three marginal relationships. Fig. 4a illustrates the extensive validation strategies that were undertaken.

**Table 1.**
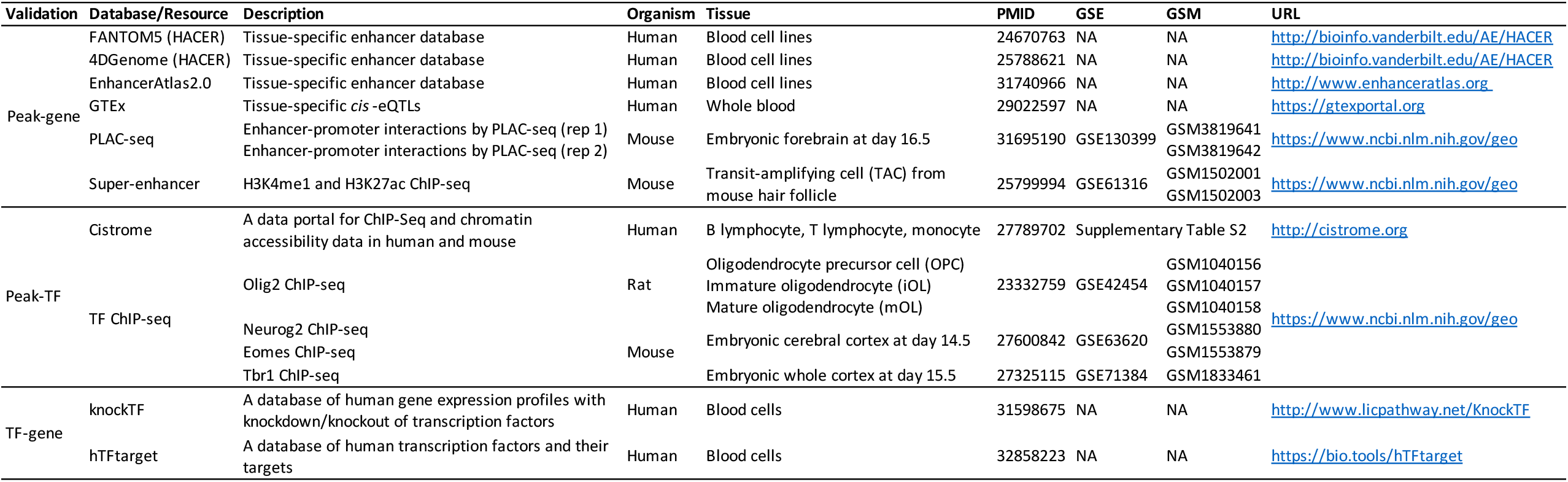
Resources for validating peak-TF-gene regulatory relationship. While there is no existing experimental approach to validate all three factors in a trio at high resolution with high throughput, we resort to existing databases and orthogonal sequencing data to validate peak-gene, peak-TF, and TF-gene pairs, completing the loop.

**Fig. 4.**
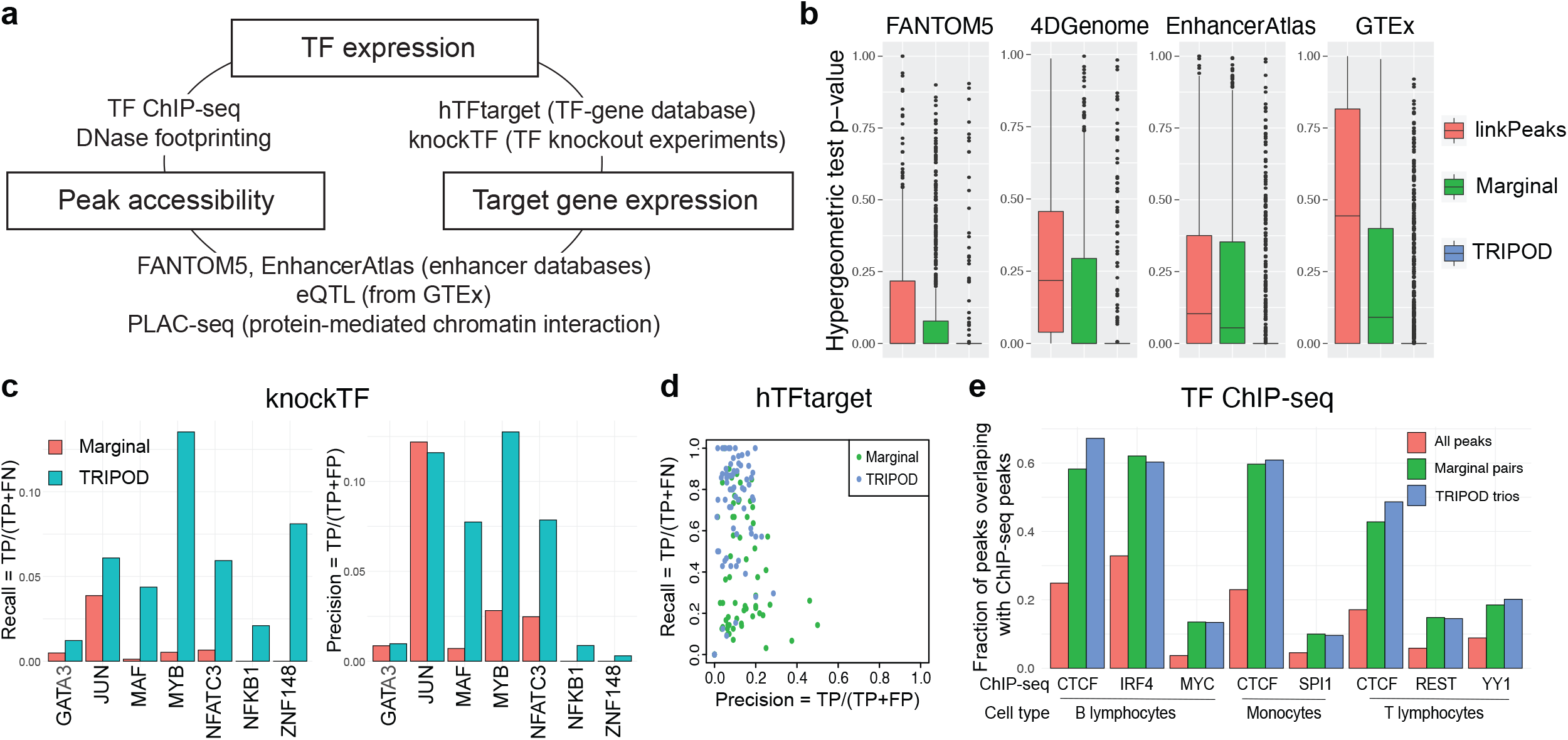
TRIPOD identified trio regulatory relationships in PBMC single-cell multiomic dataset supported by extensive validations. **a**, A schematic of validation strategies. Shown are external datasets and databases used to validate the links between peak accessibility and target gene expression (peak-gene validation), those between peak accessibility and TF expression (peak-TF validation), and those between TF expression and target gene expression (TF-gene validation). **b**, Peak-gene validation based on enhancer databases (FANTOM5, 4DGenome, and EnhancerAtlas) and tissue-specific *cis*-eQTL data from the GTEx Consortium. Box plots show distributions of *p*-values from gene-specific hypergeometric tests. **c**, TF-gene validation based on lists of TF-gene pairs from the knockTF database. **d**, Precision and recall rates for TF-gene pairs using ground truths from the hTFtarget database. **e**, Peak-TF validation based on eight cell-type-specific TF ChIP-seq datasets (B lymphocytes, monocytes, and T lymphocytes). Fractions of significantly linked peaks and all peaks that overlap with the ChIP-seq peaks are shown.

First, to validate the *cis*-linkage between peak region and target gene, we used the enhancer databases of blood and non-cancerous cells from FANTOM5^24^ (from HACER^25^), 4DGenome^26^ (from HACER^25^), and EnhancerAtlas 2.0^27^, as well as *cis*-eQTLs in the whole blood reported by the GTEx consortium^9^. We collapsed TRIPOD’s trio calls into peak-gene relationships and benchmarked against Signac’s LinkPeaks^12^ on single cells and marginal association testing on metacells; for each target gene, we performed a hypergeometric test for enrichment of the peak-gene linkages in the regulatory databases and annotations (see Methods for details). For all four databases, TRIPOD’s *p*-values for enrichment are substantially significant (Fig. 4b). When stratified by the different levels of testing, TRIPOD’s level 1 and level 2 conditional testing returns more significant enrichment compared to linkPeaks and marginal associations; the most significant enrichment is from level 1 testing matching by TF expression, which is expected since the “gold-standard” peak-gene relationship is directly captured by the level 1 testing without TF interaction (Supplementary Fig. 14a). Additionally, the unique sets of trio regulatory relationships identified by TRIPOD but not by random matching (which results in only marginally associated linkages) have significant enrichment, demonstrating the effectiveness of TRIPOD in identifying true trio relationships that complement existing methods based on marginal association testing (Supplementary Fig. 14b).

Second, to validate the TF-gene edge in the TRIPOD-identified trios, we referred to knockTF^7^, a TF knockdown/knockout gene expression database, and hTFtarget^28^, a database of known TF regulatory targets. Specifically, in knockTF, we found seven TF knockdown/knockout RNA-seq experiments in the peripheral blood category. For these TFs, we identified significantly linked genes by marginal association and by TRIPOD and found TRIPOD’s results to have significantly higher precision and recall (Fig. 4c); the improvement is robust to varying FDR thresholds (Supplementary Table 3). For hTFtarget, we obtained, for each highly variable gene, its blood-specific TFs, and calculated the gene-specific precision-recall rates – TRIPOD is more sensitive compared to marginal association testing, although both suffered from inflated “false positives,” which can also be due to the low sensitivity in the *in silico* calls by hTFtarget (Fig. 4d). Precision and recall rates with varying significance levels further confirm that TRIPOD has better agreement with existing TF knockdown/knockout data, in comparison to marginal association testing (Supplementary Fig. 15).

Third, to validate the TF-peak edge representing TF binding to peak regions, in addition to the DNA footprinting analysis shown in Supplementary Fig. 7e, we downloaded from the Cistrome portal^29^ non-cancerous ChIP-seq data from sorted human blood cells (B lymphocyte, T lymphocyte, and monocyte (Supplementary Table 4). The peaks identified by TRIPOD had a substantially higher percentage of overlap with the ChIP-seq peaks compared to the genome-wide baseline; TRIPOD’s performance is better than or on par with that from testing of marginal associations (Fig. 4e). Since ChIP-seq peaks reflect only TF binding, without consideration for the gene target of regulation, it is expected that it agrees well with marginal association test results, which are capturing such a universal relationship.

In summary, existing databases and public data of different types from a wide range of studies extensively support each of the three pairwise links in the trios reported by TRIPOD, demonstrating its effectiveness in uncovering true regulatory relationships.

### TRIPOD identifies known and novel putative regulatory relationships during mouse embryonic brain development

We next applied TRIPOD to single-cell multiomic data of 5k mouse embryonic brain cells at day 18 by 10X Genomics. The cell type labels were transferred from an independent scRNA-seq reference^30^ using SAVERCAT^31^. We kept 3,962 cells that had consistent transferred labels from seven major cell types: radial glia, neuroblast, GABAergic neuron, glutamatergic neuron, glioblast, oligodendrocyte, and Cajal−Retzius neuron (Supplementary Fig. 1b). We applied TRIPOD to 633 TFs, 1000 highly variable genes, and ATAC peaks 200kb up/downstream of the genes’ TSSs.

On the genome-wide scale, the union of TRIPOD’s level 1 and 2 tests gave a larger number of unique peak-gene pairs and TF-gene pairs than LinkPeaks^12^ and marginal association testing, respectively (Supplementary Fig. 16a). To evaluate these results, we first examined whether the peak-gene links were enriched in previously reported enhancer-promoter chromatin contacts using PLAC-seq data of mouse fetal brain^32^ (Table 1, Supplementary Fig. 16b). We observed that the regulatory links detected by both marginal association and TRIPOD showed significant enrichment in PLAC-seq contacts (Supplementary Fig. 16b). Importantly, TRIPOD detected sets of peak-gene pairs from trio relationships that were overlapping but distinct from the sets obtained by marginal association, and a substantial fraction of the links identified by TRIPOD but not by the marginal method were validated by PLAC-seq (Fig. 5a; Supplementary Fig. 16c). This suggests that TRIPOD identifies real regulatory relationships that complement those detected by existing methods.

**Fig. 5.**
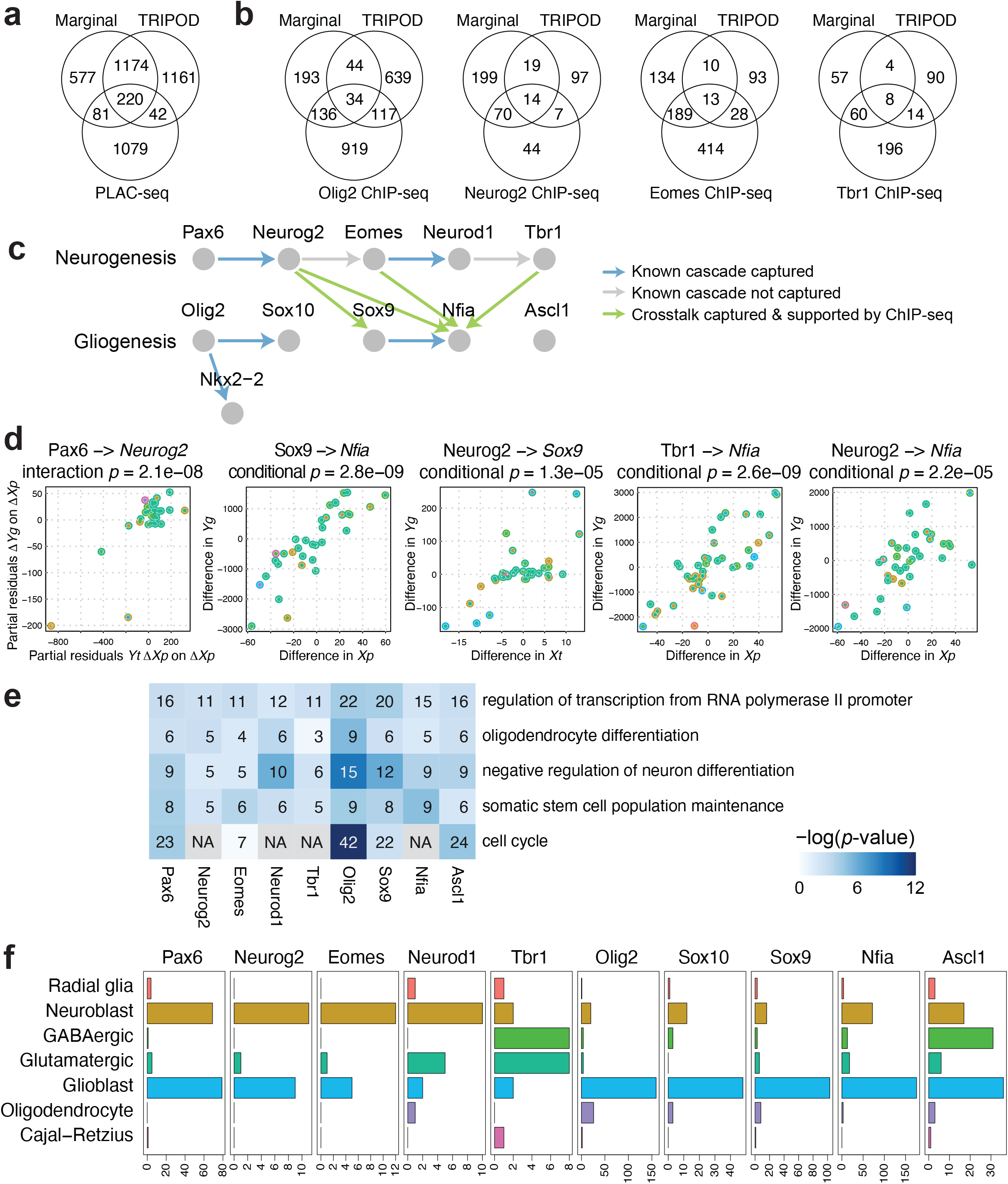
TRIPOD identified known and novel regulatory relationships during mouse embryonic brain development. **a**, Venn diagram of the number of peak-gene pairs captured by PLAC-seq, the marginal model, and the union set of TRIPOD’s level 1 and level 2 testing matching TF expression and peak accessibility. **b**, The same as **a** but for Peak-TF validation by ChIP-seq data for Olig2, Neurog2, Eomes, and Tbr1. **c**, A schematic of well-characterized TF regulatory cascades during neurogenesis and gliogenesis. **d**, Trio examples from known regulatory relationships, as well as from crosstalks supported by ChIP-seq data, captured by TRIPOD. **e**, GO analysis of putative target genes of the neurogenesis and gliogenesis TFs. The number of TRIPOD-identified target genes in the GO categories is shown. The background heatmap shows negative log *p*-values (FDR < 0.05) from hypergeometric tests examining enrichment of GO terms. **f**, Bar plots showing the number of putative cell-type-specific trios mediated by the neurogenesis- and gliogenesis-specific TFs.

We also note that the type of evidence that supports a regulatory relationship matters when compared to other types of experimental data. For example, PLAC-seq measures, for a fixed TF, the degree of promoter contacts in the TF-binding domains. Conceptually, the closest analog to this measurement in our model is level 1 association, conditioned on TF expression, between the motif-containing peak region and target gene expression. Thus, it is not surprising that this level 1 test matching by TF gives the most significant enrichment (Supplementary Fig. 16b, 17a, 18a). However, detection by TRIPOD is pre-conditioned on the expression of the target gene at a high enough level, which is irrelevant to the PLAC-seq data. Thus, not all detections made by PLAC-seq are expected to be found by TRIPOD.

To validate the links between TFs and peaks, we used publicly available ChIP-seq data for Olig2^33^, Neurog2^34^, Eomes^34^, and Tbr1^35^, TFs that play key roles in embryonic brain development (Table 1). The Olig2 ChIP-seq data were generated in three types of rat cells, oligodendrocyte precursor cells (OPC), immature oligodendrocytes (iOL), and mature oligodendrocytes (mOL), while the Neurog2, Eomes, and Tbr1 ChIP-seq data were generated in mouse embryonic cerebral cortices (see Methods for details). When TF expression was matched, TF binding peaks identified by TRIPOD level 1 tests were significantly enriched in the TF ChIP-seq peaks across all datasets except for the Olig2 ChIP-seq data of mature oligodendrocytes (mOL), which served as a negative control and had a substantially lower degree of enrichment (Supplementary Fig. 17, 18). Importantly, TRIPOD detected a substantial number of peak-TF pairs that were not detected through marginal associations but validated by ChIP-seq (Fig. 5b).

The validations and global benchmarking demonstrate TRIPOD’s effectiveness in finding real regulatory relationships. Next, we focused on a set of TFs known to play essential roles during mouse embryonic brain development. Specifically, we chose Pax6, Neurog2, Eomes, Neurod1, and Tbr1, major TFs mediating glutamatergic neurogenesis^36^, and Olig2, So×10, Nkx2-2, Sox9, Nfia, and Ascl1, which initiate and mediate gliogenesis^37^; the known regulatory cascades are shown in Fig. 5c. Here, the up and downstream TFs in a link are used as the TF and the target gene in TRIPOD’s analysis, respectively, and we established a link if at least one of the TRIPOD tests returned a positive coefficient estimate with FDR-adjusted *p*-values less than 0.01 for at least one trio involving the pair of the TF and the target gene. TRIPOD’s level 1 and level 2 testing successfully captured five out of the seven known regulatory links (Fig. 5c, d, Supplementary Fig. 19, 20); interestingly, TRIPOD’s results also suggest substantial crosstalk between the two cascades, where neurogenesis-specific TFs activate gliogenesis-specific TFs (Fig. 5c, d). ChIP-seq data of Neurog2, Eomes, and Tbr1 supported four of the crosstalk links: regulation of *Sox9* by Neurog2 and regulation of *Nfia* by Neurog2, Eomes, and Tbr1, respectively (Supplementary Fig. 21). These crosstalk links that were validated by ChIP-seq were also captured by conditional associations; two of them were captured by marginal associations (Supplementary Fig. 19). Thus, we think it is highly plausible that neurogenesis TFs activate gliogenesis genes at day 18 of embryonic mouse brain development, which is exactly when the switch is being made from neurogenesis to gliogenesis. To our best knowledge, these possible links between neurogenesis and gliogenesis pathways have not been systematically explored and thus warrant future investigation. Finally, for each of the neurogenesis and gliogenesis TFs, we performed a gene ontology (GO) analysis of their significantly linked target genes using DAVID^38^; the enriched terms were largely consistent with the regulatory functions of the TFs during neurogenesis and gliogenesis (Fig. 5e). Specifically, the mouse embryonic brain cells are collected during the transition phase between neurogenesis and gliogenesis, and the enriched terms contain oligodendrocyte differentiation and regulation of neuron differentiation, confirming TRIPOD’s calling results. Other terms, such as regulation of transcription and cell cycle, are enriched due to the transcriptional regulatory role of the TFs.

So far, we have taken advantage of the cross-cell-type variation to identify the trio regulatory relationships. To dissect cell-type-specific regulation, we next applied the influence analysis framework (see Methods for details) to the significant trios involving neurogenesis and gliogenesis TFs. For a given TF, the number of trios, for which a given cell type was influential (FDR < 0.01), is summarized in Fig. 5f, with details for specific example trios given in Supplementary Fig. 22. The analyses underpinned the cell types in which the transcriptional regulation was active, and, reassuringly, the neurogenesis and gliogenesis TFs have the most regulatory influence in neuroblasts and glioblasts, respectively. Additionally, Ascl1 is active in GABAergic neurons in addition to neuroblasts and glioblasts, consistent with its role as a GABAergic fate determinant^39^. Notably, the highly influential cell types that lead to the significant trios involving several neurogenesis-specific TFs include not only neuroblast but also glioblast, supporting our previous findings on the crosstalk between the two cascades. Notably, these results are unlikely due to the given TFs being overexpressed in the corresponding highly influential cell types, since the influential cell types were not the same as the cell types where the TFs were highly expressed (Fig. 5f, Supplementary Fig. 22, 23). Overall, TRIPOD allows fine characterization of cell-type- and cell-state-specific functions of the TFs during neurogenesis and gliogenesis.

Using this dataset, we further examined how varying window sizes and different resolutions/constructions of metacells affect the model fitting results; this led to the following observations. First, incorporating peaks 100kb/200kb up/downstream of genes’ TSSs leads to consistent and significant enrichment of validated gene-peak pairs by PLAC-seq and peak-TF pairs by ChIP-seq, while narrowing the window size down to 50kb decreased the degree of enrichment (Supplementary Fig. 17). Second, the validation results were robust to changes in resolutions of the metacells (Supplementary Fig. 18), since TRIPOD does not require the metacells to truly represent distinct and non-overlapping segments of the transcriptome space.

### TRIPOD infers lineage-specific regulatory relationships in differentiating mouse hair follicle cells

As a last example, we applied TRIPOD to SHARE-seq^13^ data (Supplementary Fig. 1c) of mouse hair follicle cells, consisting of four broadly defined cell types – transit-amplifying cells (TAC), inner root sheath (IRS), hair shaft, and medulla cells – along a differentiation trajectory. The cell-type labels were curated based on marker genes, TF motifs, and ATAC peaks from the original publication^13^; pseudotime was inferred using Palantir^40^ and overlaid on the cisTopic^41^ reduced dimensions of the ATAC domain. Cells were partitioned using both the pseudotime and the UMAP coordinates to construct metacells (Fig. 6a). Due to the low RNA coverage (Supplementary Fig. 4), we focused on 222 highly-expressed TFs, 794 highly expressed genes reported to have more than ten linked *cis*-regulatory peaks^13^, and peaks 100kb up/downstream of the genes’ TSSs.

**Fig. 6.**
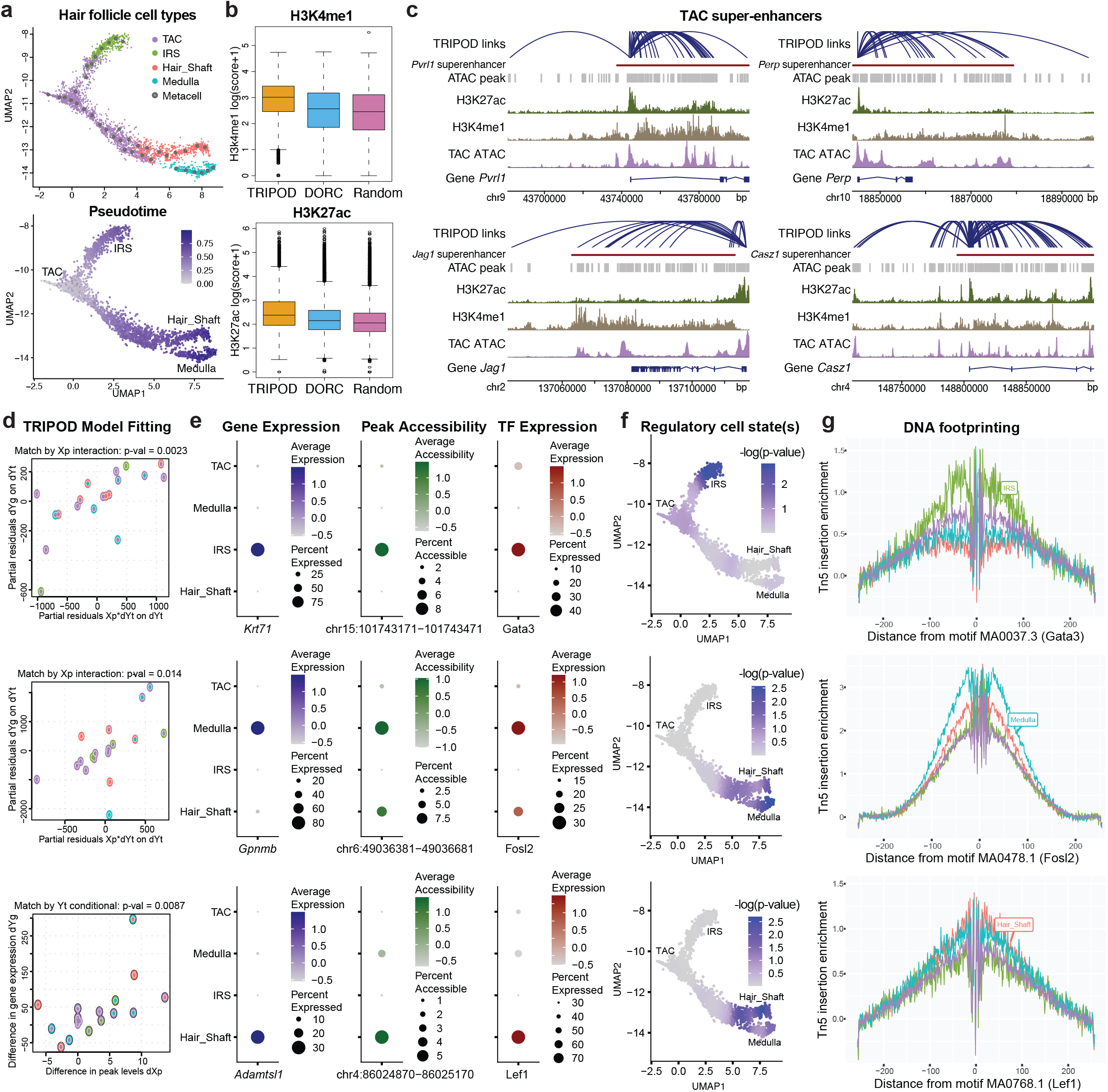
TRIPOD identified regulatory relationships in mouse hair follicles with transient cell states. **a**, UMAP embedding of hair follicle cells from the mouse skin data. Cells are colored by cell types (TAC, IRS, hair shaft, and medulla) and pseudotime. **b**, H3K4me1 and H3K27ac ChIP-seq scores for linked peaks identified by TRIPOD, DORCs (regulatory domains identified by gene-peak correlations), and randomly sampled peaks. **c**, TRIPOD’s linked peaks for four representative genes were significantly enriched in previously annotated super-enhancers in the mouse TAC population. **d**, Trios detected by TRIPOD that were active in IRS (top), medulla (middle), and hair shaft (bottom), respectively. **e**, Dot plots of gene expressions, peak accessibilities, and TF expressions across different cell types. **f**, Influence analyses identified segments along the differentiation trajectory where the regulation took effect. The colors in the UMAP embedding correspond to the smoothed *p*-values from a sampling-based approach. **g**, DNA footprinting assays showed cell-type-specific enrichments of Tn5 integration events. The findings were consistent with those from the influence analyses.

For validation, we used H3K4me1 and H3K27ac ChIP-seq data from an isolated mouse TAC population^42^ (Table 1). H3K4me1 and H3K27ac are markers for poised and active enhancers, respectively, and were used to benchmark TRIPOD’s linked peaks against previously reported domains of regulatory chromatin (DORCs)^13^, as well as randomly sampled peaks. The linked peaks by TRIPOD had higher scores for both H3K4me1 and H3K27ac than DORCs, the latter identified through marginal associations (Fig. 6b). To further validate the regulatory effects of the linked peaks, we obtained previously characterized super-enhancers (SEs) in mouse TACs^42^. Target genes of the 381 SEs were assigned based on the gene’s proximity to the SE, as well as the correlation between loss of the SE and loss of the gene transcription^42^. TRIPOD was able to successfully recapitulate the SE regions for the genes considered, with four examples shown in Fig. 6c, where significantly linked peaks mostly resided in the SEs.

To demonstrate, Fig. 6d shows regulatory trios that are specific to the IRS lineage, the hair shaft lineage, and the medulla lineage (Supplementary Fig. 24). These trios also showed significant pairwise marginal associations (Fig. 6e), lending confidence that they are real. The cell types where the regulation happens were identified by influence analysis, for which the *p*-values were smoothed along the differentiation trajectory and overlaid on the UMAP embedding (Fig. 6f). DNA footprinting analyses surveyed the enrichment of Tn5 integration events surrounding the corresponding motif sites and showed cell-type-specific enrichment (Fig. 6g), corroborating TRIPOD’s results.

## Discussion

We have considered the detection of regulatory trios, consisting of a TF binding to a regulatory region to activate or repress the transcription of a nearby gene, using single-cell RNA and ATAC multiomic sequencing data. The presented method, TRIPOD, is a new nonparametric approach that goes beyond marginal relationships to detect conditional associations and interactions on peak-TF-gene trios. We applied TRIPOD to three single-cell multiomic datasets from different species and protocols with extensive validations and benchmarks. We started our analyses with predicting gene expression from both peak accessibility and TF expression. Supervised frameworks have been proposed to predict gene expression from DNA accessibility^43^, and vice versa^44^, using matched bulk transcriptomic and epigenomic sequencing data. Blatti *et al*.^45^ showed that joint analysis of DNA accessibility, gene expression, and TF motif binding specificity allows reasonably good prediction of TF binding as measured by ChIP-seq. However, none of these methods incorporate TF expression. By selecting peaks near the genes’ TSSs and TFs with high motif scores in the selected peaks, we constructed biologically meaningful peak-TF pairs as predictors and showed that such a mechanistic model significantly boosts the prediction accuracy of gene expression.

We next considered the detection and significance assessment for individual peak-TF-gene trios, comprehensively comparing our detections with those made by tissue- and cell-type-matched PLAC-seq and ChIP-seq experiments, by *cis*-eQTL and TF knockdown/knockout studies, and by those recorded in the main enhancer databases. The comparisons show that TRIPOD detections are substantially enriched for overlap with all of these experiments, and in most cases, improve upon the overlap achieved by existing methods. It is important to note that the recall rates in the comparisons to these experiments should only be interpreted as relative metrics and not as absolute measures of sensitivity. That is because each experiment measures a biological relationship that is associated but different from what we aim to recover from TRIPOD. For example, ChIP-seq aims to capture all locations where the TF binds, regardless of which gene it is affecting, while TRIPOD aims to recover specific TF, enhancer, target gene trios. KnockTF and hTFtarget, on the other hand, aims to identify all genes whose expressions change when a TF is knocked out/down, which may not be genes that the TF directly regulates through binding. An experiment that perhaps comes closest to measuring what TRIPOD detects is PLAC-seq, which quantifies chromatin contacts anchored at genomic regions bound by specific proteins. In addition to ChIP-seq, we used PLAC-seq data to corroborate TRIPOD detections for the embryonic mouse brain data in Fig. 5a, Supplementary Fig. 16b, 17a, 18a. Here, the overlap is also far from 100%, as TRIPOD can only detect a PLAC-seq relationship if the expression of the target gene is high enough. Also, PLAC-seq cannot detect TRIPOD relationships unless the *cis*-region in question comes into direct contact with the promoter, which is not the only mechanism of gene regulation. For example, TF binding may change the local chromatin conformation as an insulator or may help recruit the binding of other TFs. Thus, it is expected that TRIPOD only recovers a small fraction of the signals identified by these experiments. For this reason, we choose to use the word “recall” rather than “sensitivity,” as we are using it as a metric of enrichment rather than as a measure of true positive rate.

Our current study is limited in several ways. A study in *Drosophila*^46^ modeled motif binding specificities and chromatin accessibilities in bulk RNA and ATAC sequencing data to predict the cooperative binding of pairs of TFs, using *in vitro* protein-protein binding experiments for validation. The detection of synergies between multiple TFs and peaks on the genome-wide scale and in a cell-type-specific manner needs further investigation. Additionally, while we have not differentiated between positive and negative regulation, TRIPOD reports both types of relationships and categorizes them by sign. While we describe the trios with a positive sign to be enhancers, it is not clear how to interpret the trios with negative signs, the latter having lower overlap with other benchmarking datasets. Transcription activation and repression have been active research areas in biology, with a lot yet unknown^47^. TRIPOD’s results provide potential targets for experimental follow-up and detailed characterization.

TRIPOD uses cell matching as a nonparametric method of computing conditional associations. One could, conceptually, match on more cell-level attributes in addition to transcription factor expression or peak level accessibility. For example, to recover true causal relationships, it seems tempting to match on more potential confounders, such as cell type. However, one should be careful in matching by additional covariates such as inferred cell type labels, as this could also reduce the signal. For example, condition-specific regulation signals that are shared across multiple (but not all) cell types would be much reduced if we were to match on cell type. For specificity, TRIPOD relies on the careful curation of inputs to the regression (using only peaks that contain the TF motif and are close to the target gene), rather than matching on all possible confounders.

Our analysis focused on three datasets where the RNA and ATAC modalities have sufficient depths of coverage. For the SHARE-seq data, the sequencing depth for RNA is very low, and thus we focused only on highly expressed genes and TFs (Fig. 6). For SNARE-seq data, whose coverage in both modalities is even lower, we focused on prediction models and not trio detection, where we saw only marginal improvement beyond existing methods^20^ (Supplementary Fig. 2b). For data where the coverage is even lower, e.g., PAIRED-seq, cross-modality metacells could not be stably formed, making such analyses impossible (Supplementary Table 1, Supplementary Fig. 4). With rapidly increasing sequencing capacity and technological advancement, TRIPOD, applied to more cells sequenced at higher depth, can uncover novel regulatory relationships at a finer resolution. With increased data resolution and cell numbers, it would then be meaningful to explore beyond the three-way relationships characterized by TRIPOD to include higher-order models that can more realistically capture the complex regulatory relationships between enhancers, modules consisting of multiple transcription factors, and the transcription of the target gene.

## Methods

### Data input and construction of metacells

Denote *X*_*ip*_ as the peak accessibility for peak *p* (1 ≤ *p* ≤ *P*) in cell *i* (1 ≤ *i* ≤ *N*), *Y*_*ig*_ as the gene expression for gene *g* (1 ≤ *g* ≤ *G*), and *Y*_*it*_ as the TF expression for TF *t* (1 ≤ *t* ≤ *T*). The TF expression matrix is a subset of the gene expression matrix, and for single-cell multiomic data, the cell entries are matched. To mitigate the effect of ATAC sparsity^48^ and RNA expression stochasticity^49^, as a first step, TRIPOD performs cell-wise smoothing by pooling similar cells into “metacells.” This, by default, is performed using the weighted-nearest neighbor method by Seurat V4^14^ to jointly reduce dimension and identify cell clusters/states across different modalities. In practice, the metacells can also be inferred using one modality – for example, RNA may better separate the different cell types^30^, and in other cases, chromatin accessibility may prime cells for differentiation^13^. For data normalization, we use sctransform^50^ and TF-IDF^11^ for scRNA-seq and scATAC-seq, respectively, followed by dimension reduction and clustering^12^. To account for peaks overlapping with other genes (Supplementary Fig. 6b), TRIPOD has the option to either remove the overlapped peaks or to adjust the peak accessibilities by the expressions of the overlapped genes, in a similar fashion to MAESTRO^18^. To reconstruct the RNA and ATAC features for the metacells, we take the sum of the integer-valued ATAC and RNA read counts across cells belonging to the metacells; library size is adjusted for both the RNA and ATAC domain by dividing all counts by a metacell-specific size factor (total read counts divided by 10^6^). For the analyses presented in the manuscript, position frequency matrices (PFM) were by default obtained from the JASPAR database^51^, and we used 633 and 107 pairs of TFs and motifs annotated in human and mouse, respectively. TRIPOD provides an option to use a more comprehensive set of motif annotations from the HOCOMOCO^52^ database. TRIPOD also allows for a binding motif to be shared across multiple TFs, as well as user-defined and/or de novo motifs. We additionally examined the effects of combining the accessibilities of ATAC peaks containing the TF binding sites within the window centered at the gene’s TSS and using the combined accessibility as input; we did not observe an improvement in model performance (Supplementary Fig. 25).

### RNA prediction by TF expression and peak accessibility

To predict RNA from ATAC, Signac^12^ and Cicero^17^ take the sum of peak accessibilities in gene bodies and promoter regions to construct a pseudo-gene activity matrix: 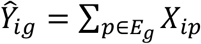, where *E*_*g*_ is the set of peaks within gene bodies and upstream regions of TSSs. Instead of directly taking the sum, MAESTRO^18^ adopts a “regulatory potential” model by taking the weighted sum of accessibilities across all nearby peaks: 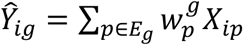, with weights 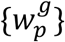 pre-calculated based on existing gene annotations. Specifically, the method weighs peaks by exponential decay from TSS, sums all peaks on the given gene exons as if they are on the TSS, normalizes the sum by total exon lengths, and excludes the peaks from promoters and exons of nearby genes. The strategy to take the unweighted/weighted sum of accessibility as a proxy for expression has been adopted to align the RNA and ATAC modalities when scRNA-seq and scATAC-seq are sequenced in parallel from the same cell population but not the same cells^11^. For single-cell multiomic data, sci-CAR^19^ performs feature selection to identify *cis*-linked peaks via a LASSO regression: 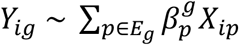, where an L1 regularization is imposed on 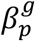. Compared to MAESTRO, which pre-fixes the weights 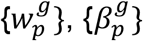 are estimated from the data by regressing RNA against matched ATAC data. What we propose is a feature selection model involving both peak accessibility and TF expression: 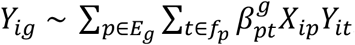, where *f*_*p*_ ontains the set of TFs with high-scoring binding motifs in peak *p* inferred from the JASPAR database^51^.

### TRIPOD model and trio regulatory relationship

For a given target gene *g*, a peak *p* within a window centered at the gene’s TSS, and a TF *t* whose binding motif is high-scoring in the peak, TRIPOD infers the relationship between a regulatory trio (*p, t, g*). TRIPOD focuses on one trio at a time and goes beyond the marginal associations to characterize the function *Y*_*g*_ = *f*(*X*_*p*_, *Y*_*t*_). In what follows, we first describe TRIPOD’s matching-based nonparametric approach and then describe a linear parametric approach, followed by a discussion on the connections and contrasts between the two approaches.

For each cell *i* whose TF expression is above a threshold *δ* (we only carry out testing in cells that express the TF), we carry out a minimum distance pairwise cross-match based on {*Y*_*it*_|*Y*_*it*_ > *δ*}. Let {(*i*_*j*_, *i*_*j*_*)} be the optimal matching, after throwing away those pairs that have 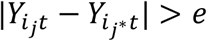. For each pair *j, i*_*j*_ and *i*_*j*_* are two metacells with matched TF expression, for which we now observe two, possibly different, values 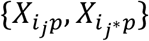 for peak *p*, as well as two corresponding values 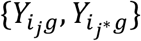 for gene *g*. We then compute the following auxiliary differentials within each pair:

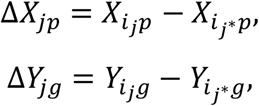

as well as

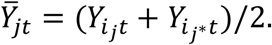

For level 1 testing of conditional association, we estimate 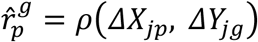, where *ρ* is Spearman correlation, and test 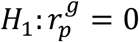. For level 2 testing of interaction, we perform a regression 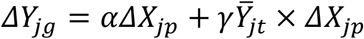, set 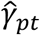 to be the least-squares solution for γ, and test *H*_2_: γ_*pt*_ = 0. For visualization of the model fitting, we take the partial residuals of Δ*Y*_*jg*_ and 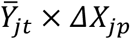 on Δ*X*_*jp*_, respectively. Note that even though TF expression is not included in the model as a main term, it is controlled for (and not just in the linear sense) by the matching. Similarly, we can also perform this procedure matching by peak accessibility. As a summary, for level 1 testing of conditional association, we have:

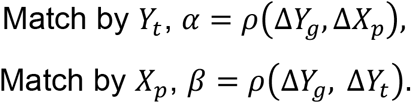

For level 2 testing of (TF expression)×(peak accessibility) interaction effects, we have:

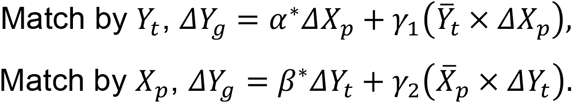

To test for the conditional associations and interactions, we can also apply a parametric method, such as multiple linear regression:

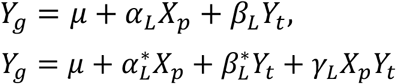

See Supplementary Fig. 26 for linear testing results for trios shown in Fig. 3 and Fig. 6. The estimated coefficients from the nonparametric and parametric methods are correlated on the global scale (Supplementary Fig. 13, 27), and their interpretations are similar: α and α_*L*_ estimate the change in gene expression per change in peak accessibility, fixing TF expression; *β* and *β*_*L*_ estimate the change in gene expression per change in TF expression, fixing peak accessibility; γ_1_ and γ_*L*_ measure how the change in gene expression per change in peak accessibility at each fixed TF expression relies on the TF expression; γ_2_ and γ_*L*_ measure how the change in gene expression per change in TF expression at each fixed peak accessibility relies on the peak accessibility. However, the underlying models and assumptions are different. Matching controls for not just the linear variation in the matched variable, but also any nonlinear variation. This contrasts with adding the variable as a covariate in the linear regression, where we simply remove linear dependence. The main motivation for using the matching model above is our reluctance to assume the simple linear relationship. Additionally, we use the rank-based Spearman correlation, which will not be driven by outliers – a “bulk” association between ranks is needed for significance. Thus, the nonparametric model of TRIPOD is more stringent (Supplementary Fig. 28) and more robust to outliers.

### Identifying regulatory cell type(s) and cell state(s)

For the significant trios detected by TRIPOD, we next seek to identify the underlying regulatory cell type(s). Specifically, we carry out a cell-type-specific influence analysis to identify cell types that are highly influential in driving the significance of the trio. Traditional approaches (e.g., the Cook’s distance and the DFFITs) delete observations one at a time, refit the model on remaining observations, and measure the difference in the predicted value from the full model and that from when the point is left out. While they can be readily applied to detect “influential” metacells one at a time (Supplementary Fig. 7a,b), these methods do not adjust for the degree of freedom properly when deleting different numbers of metacells from different cell types. That is, they do not account for the different numbers of observations that are simultaneously deleted. Additionally, both methods adopt a thresholding approach to determine significance, without returning *p*-values that are necessary for multiple testing correction. We, therefore, develop a sampling-based approach to directly test for the influence of multiple metacells and to return *p*-values (Supplementary Fig. 7c).

Here, we focus on the linear model for its ease of computation: 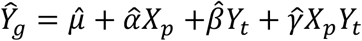. Given a set of observations *I* = {*i*: *i*th metacell belongs to a cell type}, we remove these metacells, fit the regression model, and make predictions: 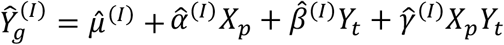. The test statistics are the difference in the fitted gene expressions 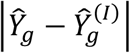. We generate the null distribution via sampling. Specifically, within each sampling iteration, we sample without replacement the same number of metacells, denoted as a set of *I*^*^, delete these observations, and refit the regression model on the remaining observations: 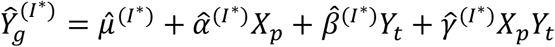. The *p* -value is computed across *K* sampling iterations as 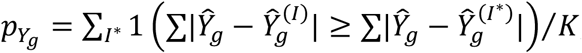, where 1() is the indicator function. In addition to testing each cell type separately, the framework can be extended to test for the influence of cell-type groups. For example, in Fig. 3, we reconstruct the cell-type hierarchy using expression levels of highly variable genes from the RNA domain and carry out the aforementioned testing scheme at each split for its descendent cell types in the hierarchical structure.

For transient cell states, TRIPOD first identifies the neighbors of each metacell along the trajectory and then carries out metacell-specific testing by simultaneously removing each metacell and its neighbors using the framework described above. The resulting *p*-values are, therefore, smoothed and can be visualized in the UMAP plot (Fig. 6f and Supplementary Fig. 22) to identify the underlying branches/segments that are key in defining the significant regulatory trio. This approach can be directly applied to cells with branching dynamics without the need to isolate cell subsets or to identify cell types.

### Validation resources and strategies

Resources for validating the trio regulatory relationships are summarized in Table 1. To validate the peak-gene relationships, we referred to existing enhancer databases: FANTOM5^24^ links enhancers and genes based on enhancer RNA expression; 4DGenome^26^ links enhancers and genes based on physical interactions using chromatin-looping data including 3C, 4C, 5C, ChIA-PET, and Hi-C; EnhancerAtlas 2.0^27^ reports enhancers using 12 high-throughput experimental methods including H3K4me1/H3K27ac ChIP-seq, Dnase-seq, ATAC-seq, and GRO-seq. We only focused on blood and non-cancerous cells from these databases (Fig. 4b). A list of *cis*-eQTLs within the whole blood mapped in European-American subjects was downloaded from the GTEx consortium^9^ (Fig. 4b). For the mouse embryonic brain dataset, we additionally adopted H3K4me3-mediated PLAC-seq data^32^, which reported enhancer-promoter chromatin contacts mapped in mouse fetal forebrain (Fig. 5a, Supplementary Fig. 16b, 17a, 18a). For the mouse skin dataset, we adopted TAC-specific ChIP-seq data of H3K4me1 and H3K27ac^42^, markers for poised and active enhancers, respectively (Fig. 6b); we also obtained previously reported super-enhancers in mouse TACs from *in vivo* studies^42^ (Fig. 6c). Genomic coordinates were lifted over from mm9 to mm10 when necessary.

To validate the TF-gene relationships in the PBMC data, we utilized the knockTF^7^ and the hTFtarget^28^ databases. knockTF interrogates the changes in gene expression profiles in TF knockdown/knockout experiments to link the TFs to their target genes in a tissue- or cell-type-specific manner. We downloaded 12 experiments, corresponding to 12 TFs (BCL11A, ELK1, GATA3, JUN, MAF, MYB, NFATC3, NFKB1, STAT3, STAT6, TAL1, and ZNF148) in the peripheral blood category, and focused on seven TFs that have at least one linked gene by any model benchmarked (Fig. 4c; Supplementary Table 3). hTFtarget computationally predicts TF-gene relationships using ChIP-seq data, and we manually downloaded the TFs associated with each of the top 100 highly variable genes in the blood tissue (Fig. 4d; Supplementary Fig. 15).

For peak enrichment analysis compared to the existing enhancers, *cis*-eQTLs, and enhancer-promoter contacts, we carried out a hypergeometric test as follows. Let *k* be the number of significantly linked peaks, *q* be the number of significantly linked peaks that overlap with annotations (e.g., annotated enhancers), *m* be the number of peaks that overlap with the annotations, and *n* be the number of peaks that do not overlap with annotations. The *p*-value of enrichment is derived from the hypergeometric distribution using the cumulative distribution function, coded as phyper(q, m, n, k, lower.tail=F) in R. We used this hypothesis testing framework to validate and benchmark the reported peak-gene links, with results shown in Fig. 4b.

To validate the peak-TF relationships, we downloaded non-cancerous cell-type-specific ChIP-seq data of human blood (B lymphocyte, T lymphocyte, and monocyte) from the Cistrome^29^ portal for the PBMC data (Fig. 4e, Supplementary Table 4), and ChIP-seq data of Olig2^33^, Neurog2^34^, Eomes^34^, and Tbr1^35^ for the mouse embryonic brain data. The Olig2 ChIP-seq data were generated in three types of rat cells: data from oligodendrocyte precursor cells (OPC) and immature oligodendrocytes (iOL) were used for validation, while data from mature oligodendrocytes (mOL) serve as a negative control^33^. Genomic coordinates were converted from rn4 to mm10. The Neurog2 and Eomes ChIP-seq data were generated in mouse embryonic cerebral cortices at day 14.5^34^; the Tbr1 ChIP-seq data was generated in the whole cortex dissected from embryos at day 15.5^35^. In addition, DNA footprinting signatures were corrected for Tn5 sequence insertion bias and stratified by cell types using the Signac package^12^ and can be used to validate the identified TFs/motifs in a cell-type-specific manner (Fig. 6g, Supplementary Fig. 7e). Hypergeometric tests for peak enrichment in TF binding sites by ChIP-seq were carried out (Supplementary Fig. 17b-d, 18b-d). The results presented in Fig. 4e were obtained in several steps: (i) we obtained sets of trios, for which B cells, T cells, and monocytes were significantly influential; (ii) we applied TRIPOD and took the union set of the significant trios; and (iii) we took the intersection between the trios obtained by the two types of analyses, collapsed the trios to TF-peak relationships, and computed the fraction of peaks overlapping ChIP-seq peaks.

## Data availability

This study analyzed existing and publicly available single-cell RNA and ATAC multiomic data. 10X Genomics single-cell multiomic datasets of PBMC (10k and 3k) and mouse embryonic brain were downloaded https://support.10xgenomics.com/single-cell-multiome-atac-gex/datasets. SNARE-seq data of adult mouse brain and SHARE-seq data of mouse skin are available from the Gene Expression Omnibus (GEO) database with accession numbers GSE126074 and GSE140203. A detailed data summary is provided in Supplementary Table 1. Validation resources based on existing databases and high-throughput sequencing data are summarized in Table 1 and Supplementary Table 4.

## Code availability

TRIPOD is compiled as an open-source R package available at https://github.com/yharigaya/TRIPOD. Scripts used for analyses carried out in this paper are deposited in the GitHub repository.

## Acknowledgments

This work was supported by the National Institutes of Health (NIH) grant R35 GM133712 (to C.Z.), R01 HG006137 (to N.R.Z), R01 GM125301 (to N.R.Z.), and R35 GM138342 (to Y.J.). The authors thank Dr. Sai Ma for support and guidance on the SHARE-seq data, Manas Tiwari for help on accessing the hTFtarget database, and Drs. Yun Li, Michael Love, Li Qian, and Jason Stein for helpful discussions and comments.

## Author contributions

Y.J. and N.R.Z. initiated and envisioned the study. Y.J., Y.H., and N.R.Z. formulated the model and developed the algorithm. Y.J. led the analyses for gene expression prediction, PBMC, and mouse skin. Y.H. led the analyses for mouse embryonic brain. Z.Z. processed reference datasets and performed cell-type label transfer. H.Z. and C.Z. provided support on validation, offered consultation, and contributed to result interpretation. Y.J., Y.H., and N.R.Z. wrote the manuscript, which was read and approved by all authors.

