## Supplementary Materials for "Nonparametric Interrogation of Transcriptional Regulation in Single-Cell RNA and Chromatin Accessibility Multiomic Data"

### Supplementary Figures

**Supplementary Fig. 1 | Reduced dimensions for single-cell multiomic datasets. a-c**, UMAP[1] embedding of single cells for the 10X Genomics PBMC, 10X Genomics mouse embryonic brain, and SHARE-seq[2] mouse skin. The three columns show UMAP embedding based on RNA, ATAC, and WNN[3], respectively, with colors corresponding to transferred/inferred cell types. The last column shows the UMAP embedding overlaid with metacell assignments.

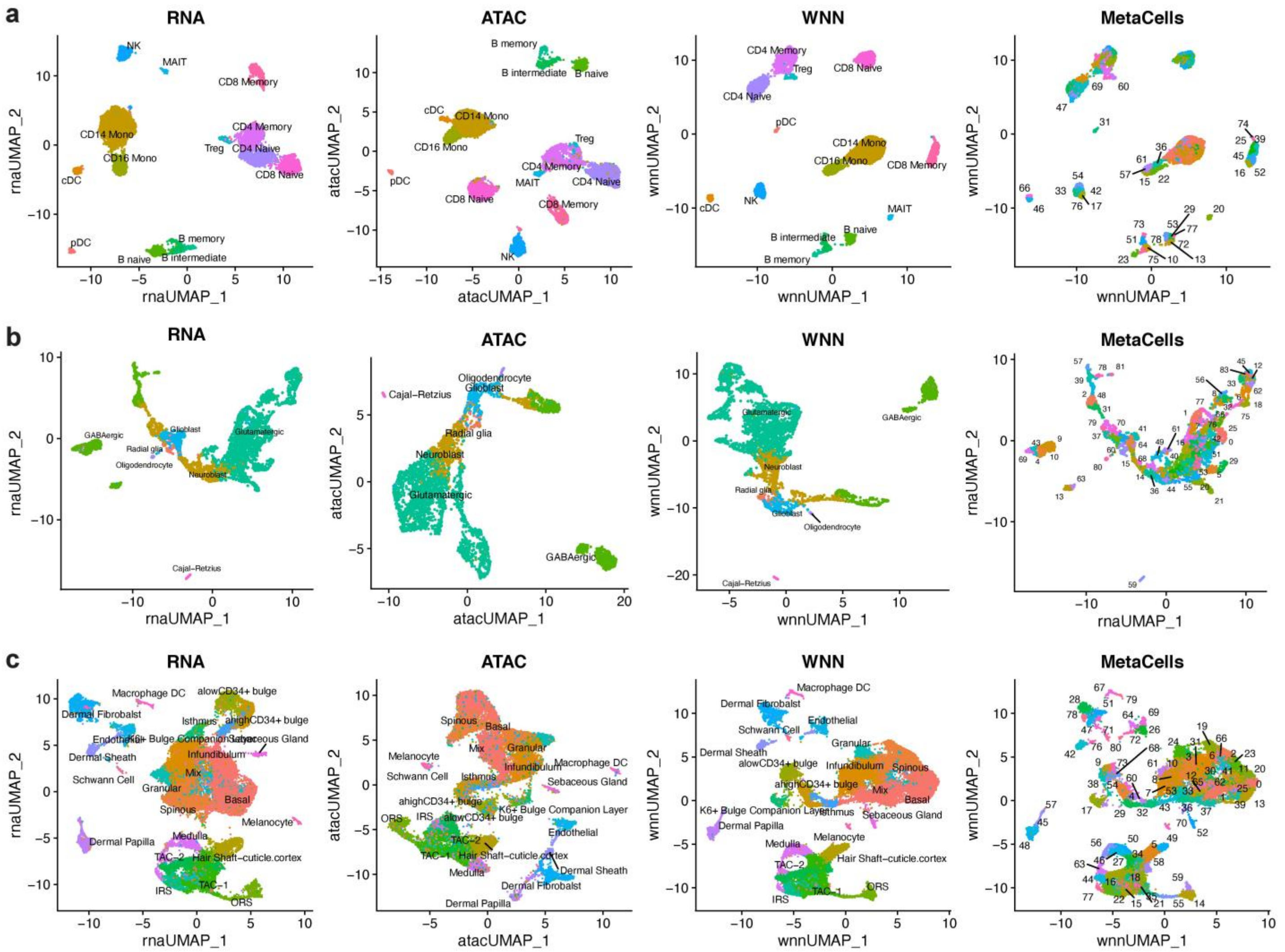

**Supplementary Fig. 2 | Reduced dimensions and RNA prediction for SNARE-seq adult mouse brain data[4]. a**, UMAP embedding of single cells based on RNA, ATAC, and WNN, respectively. **b**, Box plots showing distributions of Pearson correlations between observed and out-of-fold predicted RNA expression levels across top 1000 highly variable genes. The horizontal axis represents the sizes of the windows centered at TSSs.

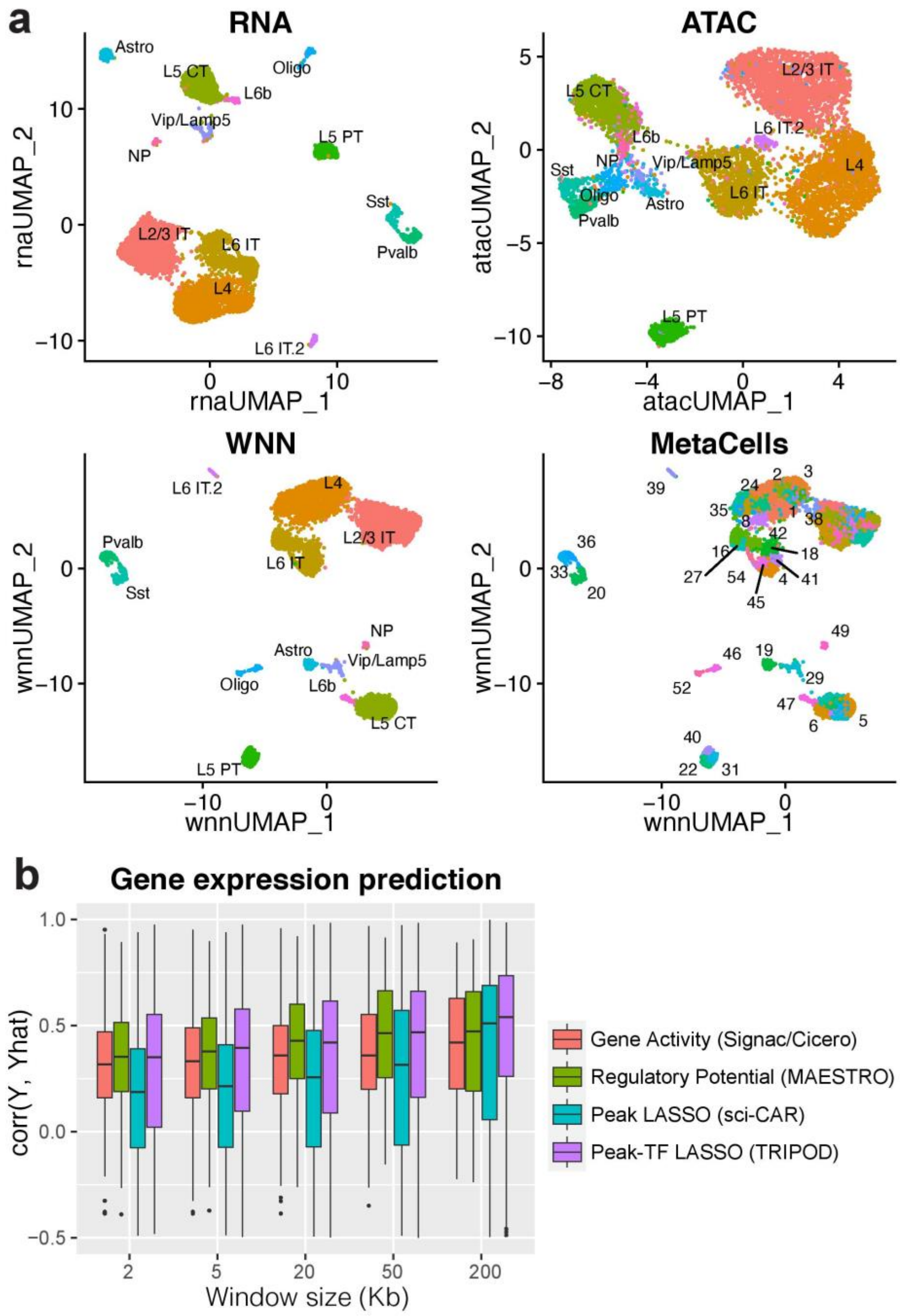


**Supplementary Fig. 3 | RNA prediction using independent training and testing datasets of PBMC.** We trained the supervised prediction model using the dataset of 10k PBMCs, adopted an independent single-cell multiomic dataset of 3k PBMCs as a testing dataset. We merged the two datasets to align genes, peaks, and TFs, and showed that the peak-TF LASSO model significantly increases the prediction accuracy. Distributions of Pearson correlations between observed and predicted RNA expression levels from the testing dataset are shown.

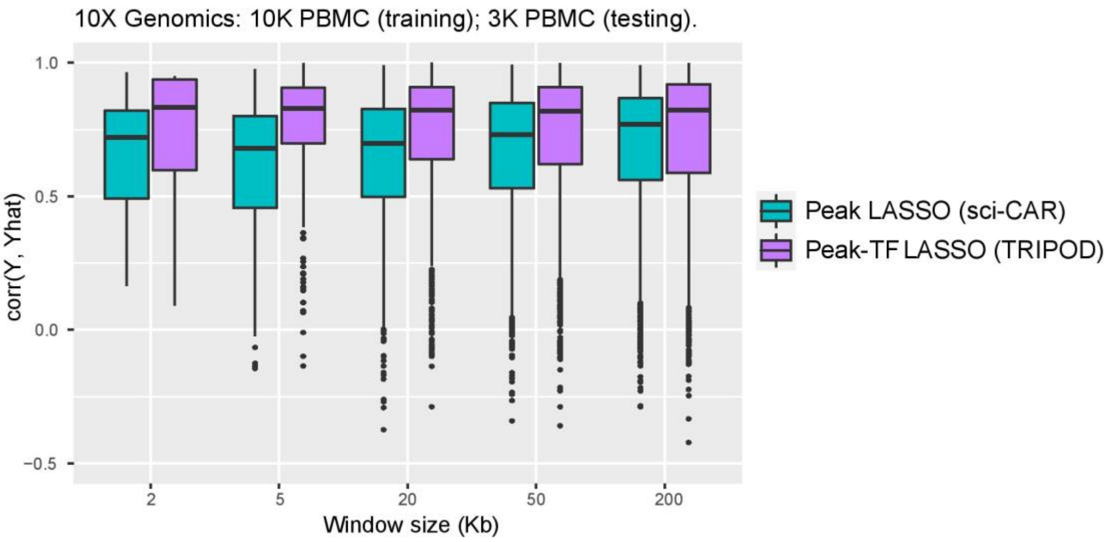

**Supplementary Fig. 4 | RNA and ATAC coverage across different single-cell multiomic sequencing protocols.** TRIPOD was applied to the 10X PBMC, 10X mouse embryonic brain, and SHARE-seq mouse skin data to detect regulatory trios. Data from SNARE-seq[4] and PAIRED-seq[5] suffer from low sequencing depth.

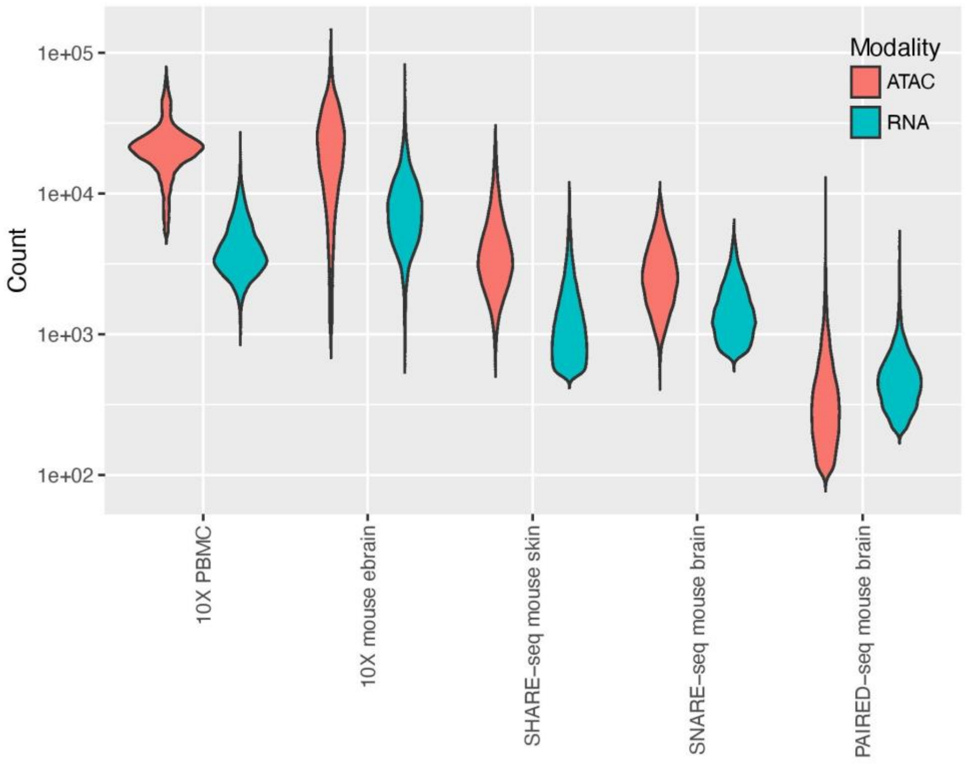

**Supplementary Fig. 5 | Visualization of target gene expression levels, chromatin accessibility in the ATAC peak regions, and TF expression levels for example trios. a-f,** The levels of gene expression (left), peak accessibility (middle), and TF expression (right) are shown on UMAP embeddings for example trios in Fig. 2c (**a**), Fig. 2d (**b**), Fig. 3a (**c**), and Fig. 3b (**d**).

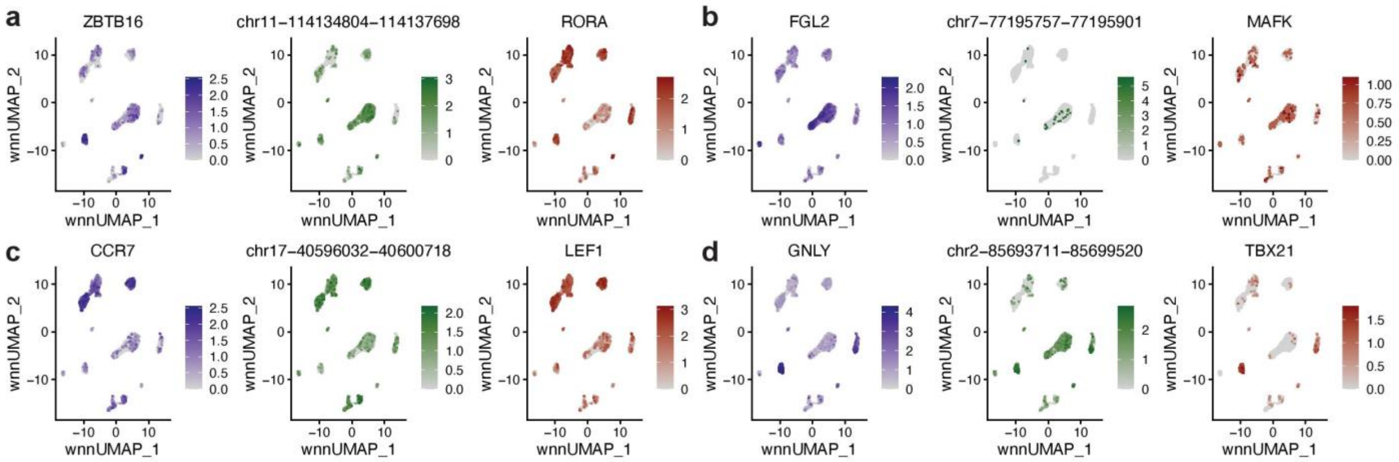

**Supplementary Fig. 6 | Summary statistics of peaks, genes, and motifs in the PBMC data. a**, Histograms of the number of peaks within regions of 100kb/200kb up and downstream of genes' TSSs. **b**, Histograms of the number of genes within regions of 100kb/200kb up and downstream of genes' TSSs. **c**, Histogram of the number of motifs per peak. Of the many possible and biologically meaningful peak-TF-gene combinations, TRIPOD proceeds to scan for trios with significant conditional associations.

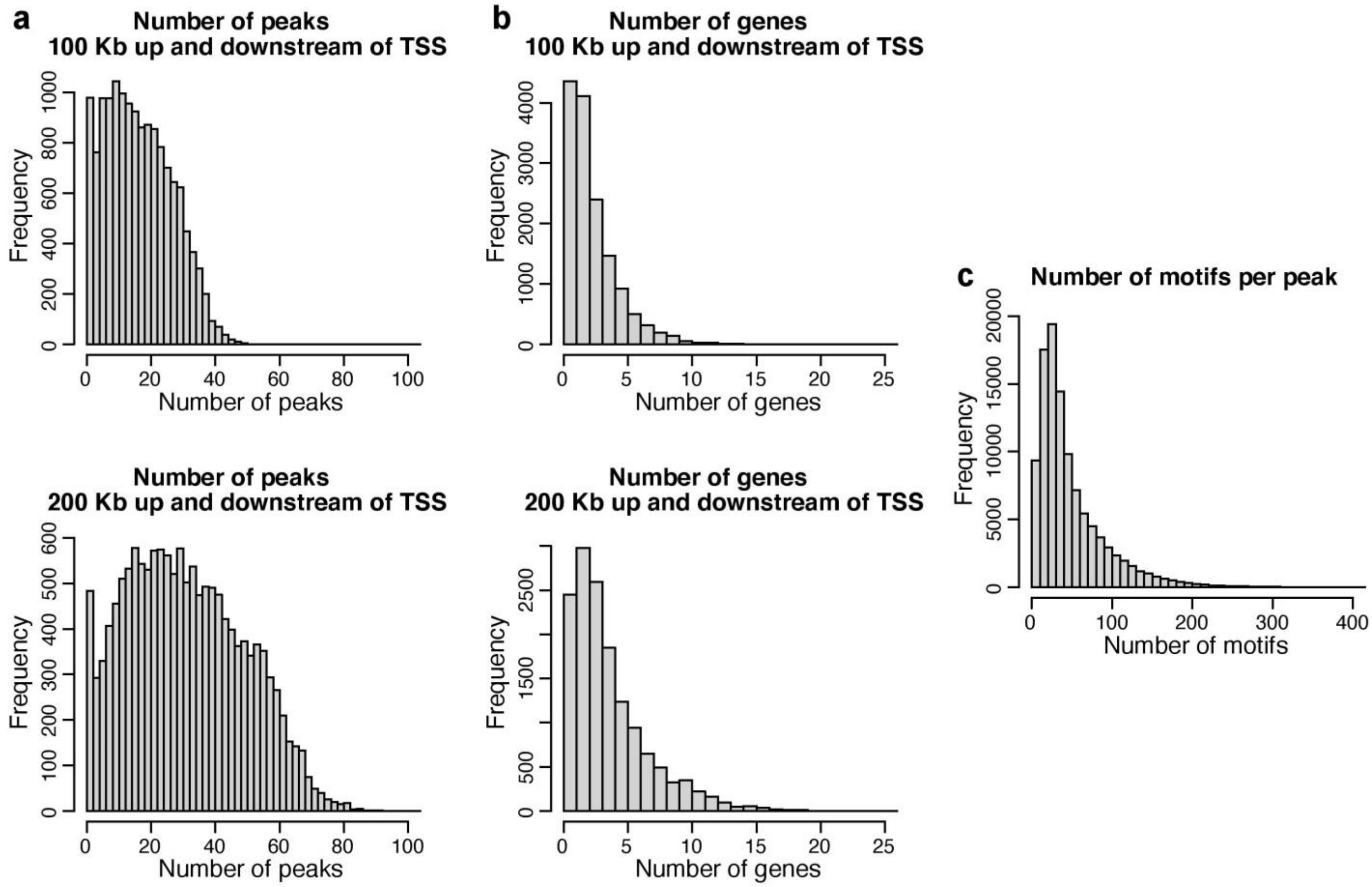

**Supplementary Fig. 7 | Identification of putative cell-type-specific trio regulatory relationships in PBMC.** Results from cell-type-specific influence analyses are shown for the same example trios as in Fig. 3. **a**, Metacell-specific Cook's distance. **b**, Metacell-specific DFFITS. **c**, Cell-type-specific influential $p$-values. **d**, Cell-type hierarchies constructed from the RNA domain using highly variable genes. Red/gray circles indicate whether removal of the corresponding branches of metacells significantly changes the model fitting; crosses indicate that removal of the groups of metacells resulted in inestimable coefficients. **e**, Cell-type-specific DNA footprinting signatures of the TF binding motifs. The enrichment results supported the key regulatory cell types identified from the influence analyses.

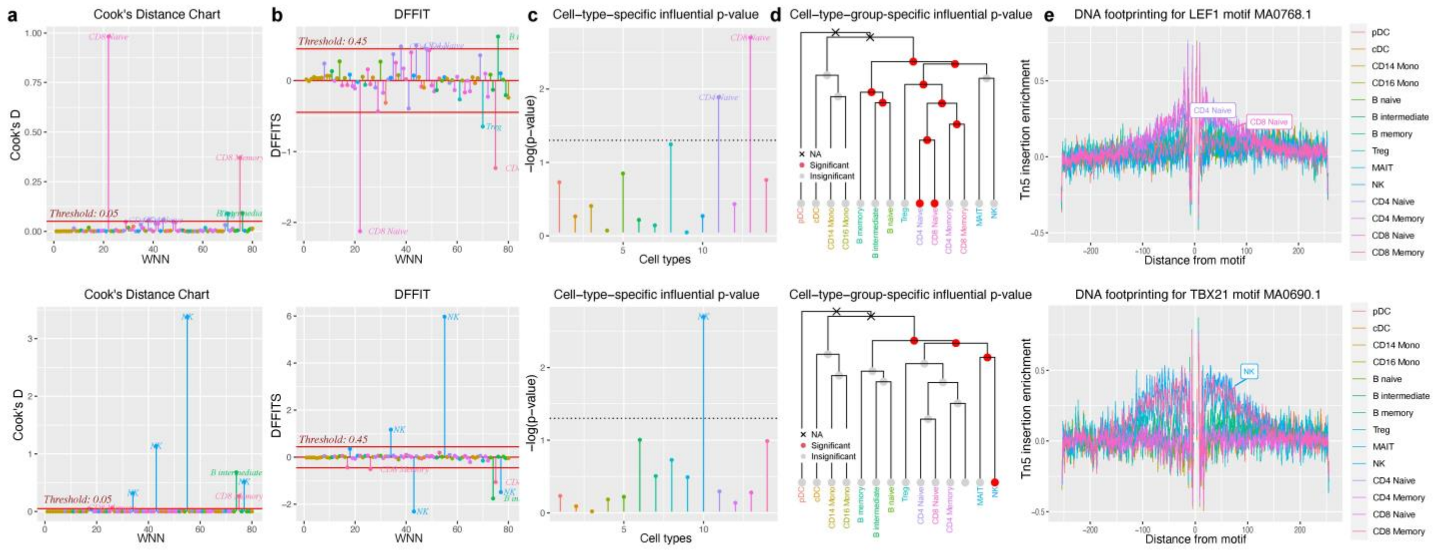

**Supplementary Fig. 8 | Regulatory relationships identified by LinkPeaks, marginal association, and TRIPOD in the PBMC data. a**, Venn diagrams of the number of peak-gene pairs captured by LinkPeaks[6], marginal association between gene expression and peak accessibility, and TRIPOD for representative target genes (*CCR7*, *GNLY*, *FCGR3A*, and *MS4A1*). For TRIPOD, the union set between level 1 and level 2 testing matching by TF expression is shown. **b**, Venn diagrams of the number of TF-gene pairs captured by marginal association between gene expression and TF expression and TRIPOD. For TRIPOD, the union set between level 1 and level 2 testing matching by peak accessibility is shown. TRIPOD complements and contrasts with existing methods based on marginal associations.

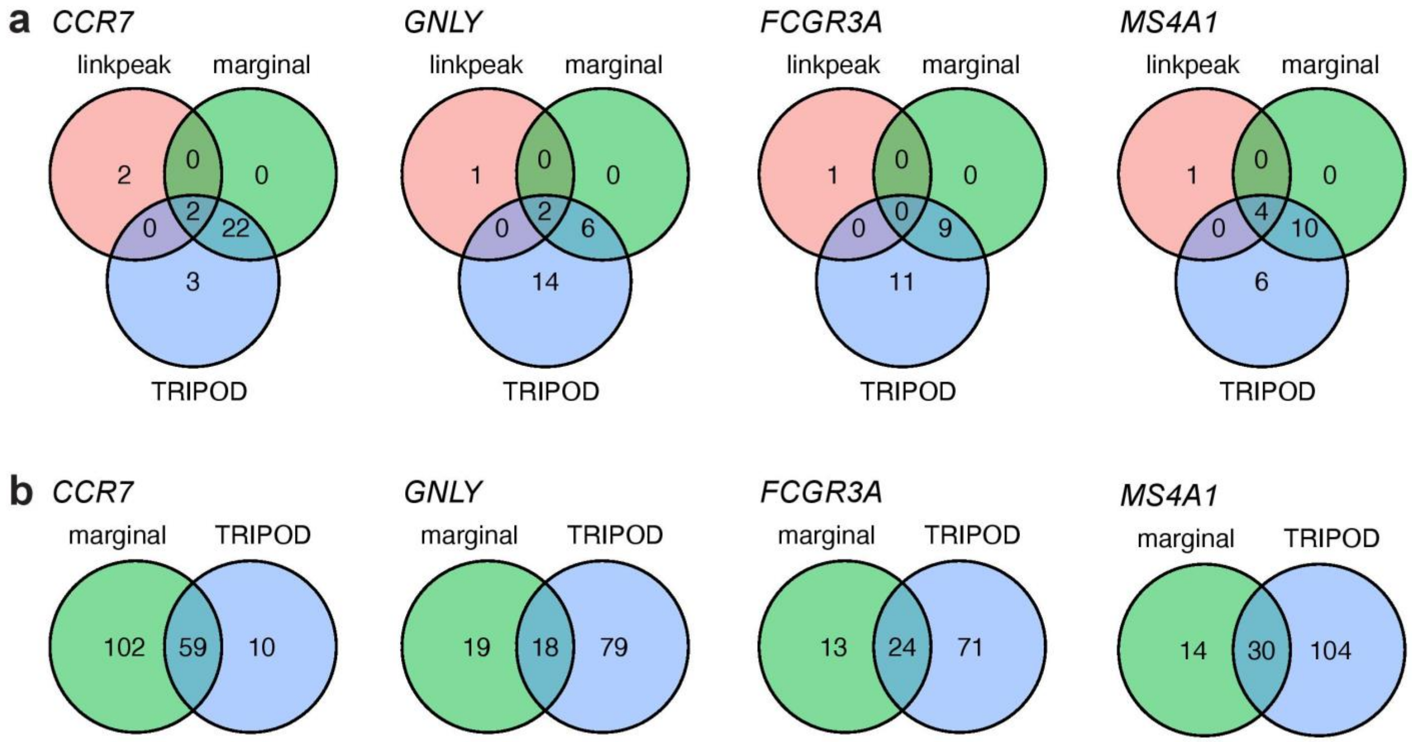

**Supplementary Fig. 9 | Level 1 and level 2 conditional testing results from TRIPOD's matching scheme and random matching.** Results from TRIPOD's matching scheme and those from random matching also overlap but exhibit substantial differences, especially for level 2 testing.

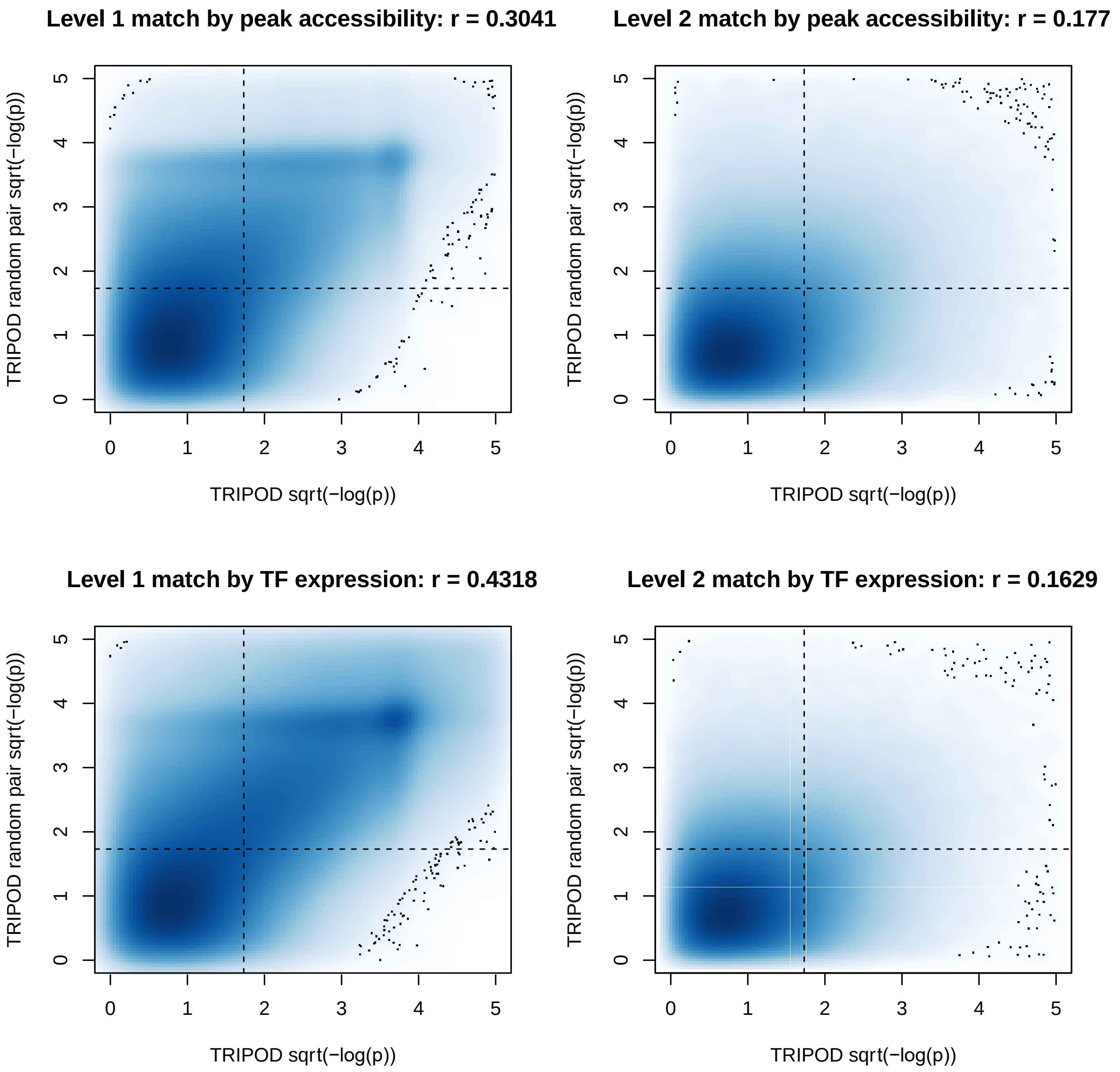

**Supplementary Fig. 10 | Gene-specific level 1 and level 2 conditional testing results from TRIPOD's matching scheme and random matching.** Results from TRIPOD's matching scheme and those from random matching also overlap but exhibit substantial differences, especially for level 2 testing.

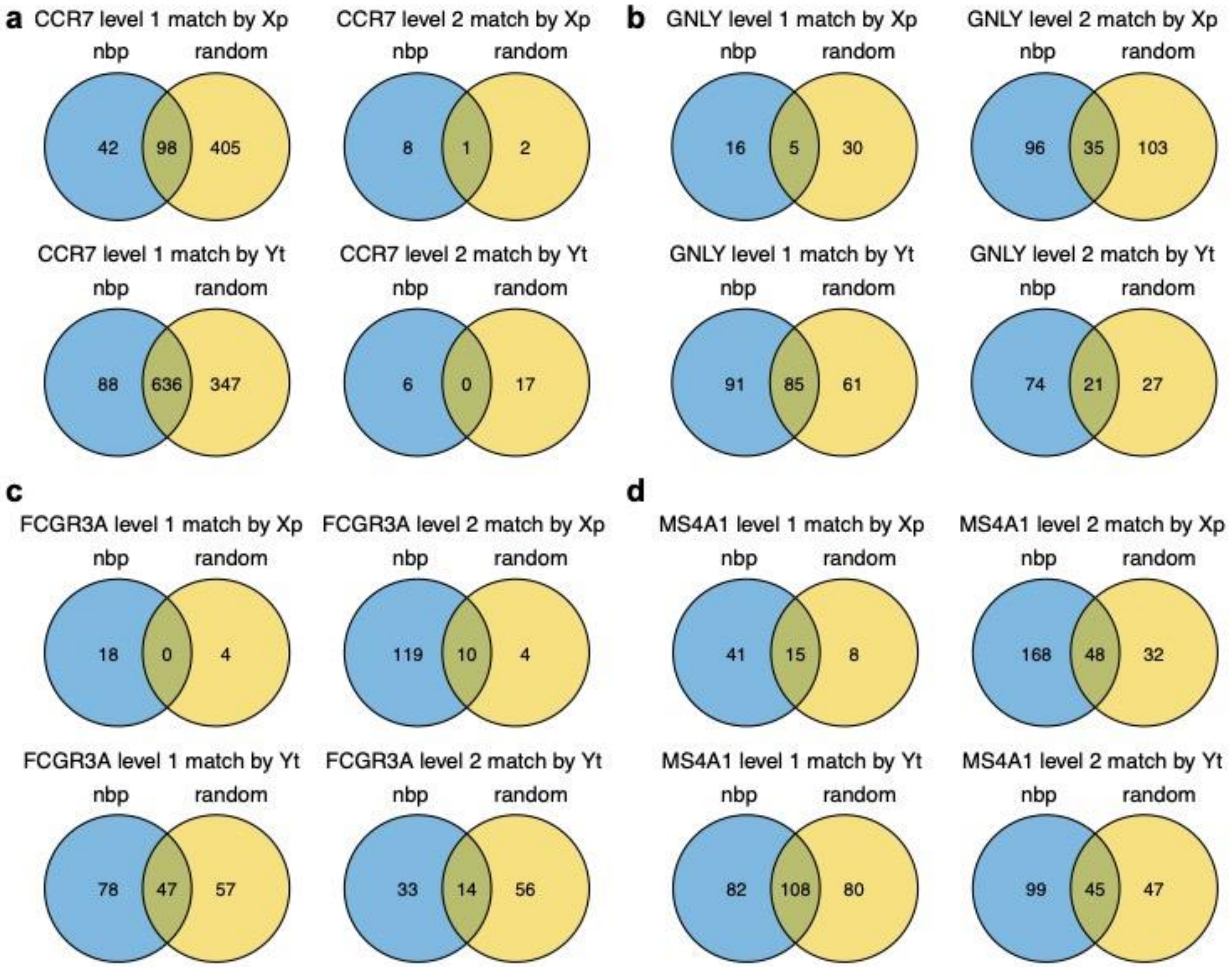

**Supplementary Fig. 11 | Random matching suffers from false positive and false negative in the two PBMC counter examples.** The same two example trios from Fig. 2 are shown with random matching, which is equivalent to marginal testing and results in false negative (**a**) and false negative (**b**).

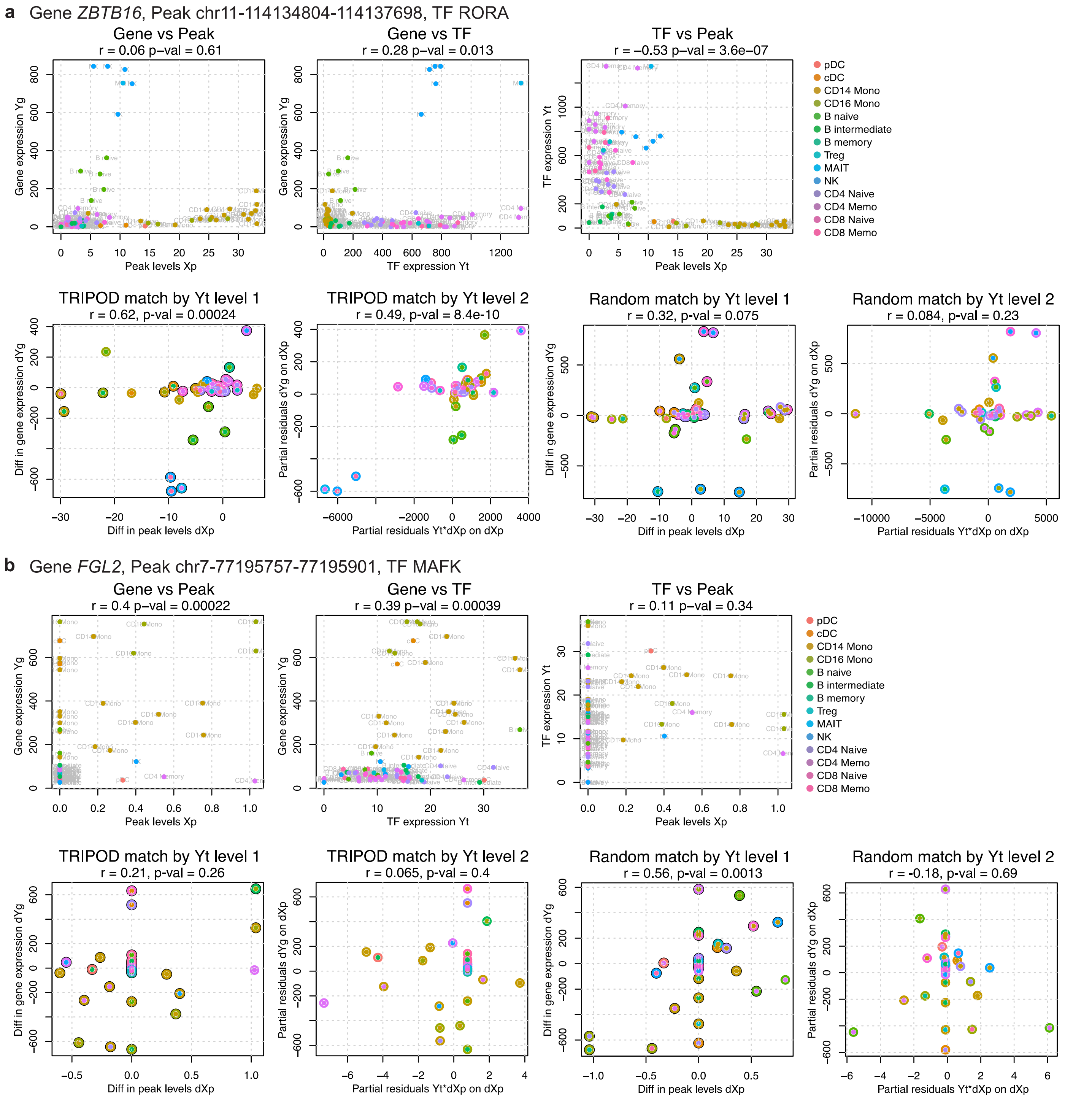

**Supplementary Fig. 12 | TRIPOD controls for type I error via permutation-based analysis. a**, TRIPOD's nominal $p$-value distributions on the PBMC multiomic data. **b**, Peak accessibilities and TF expressions are randomly permuted to generate trios under the null. TRIPOD's $p$-value distributions are uniformly distributed, indicating TRIPOD's good type I error control.

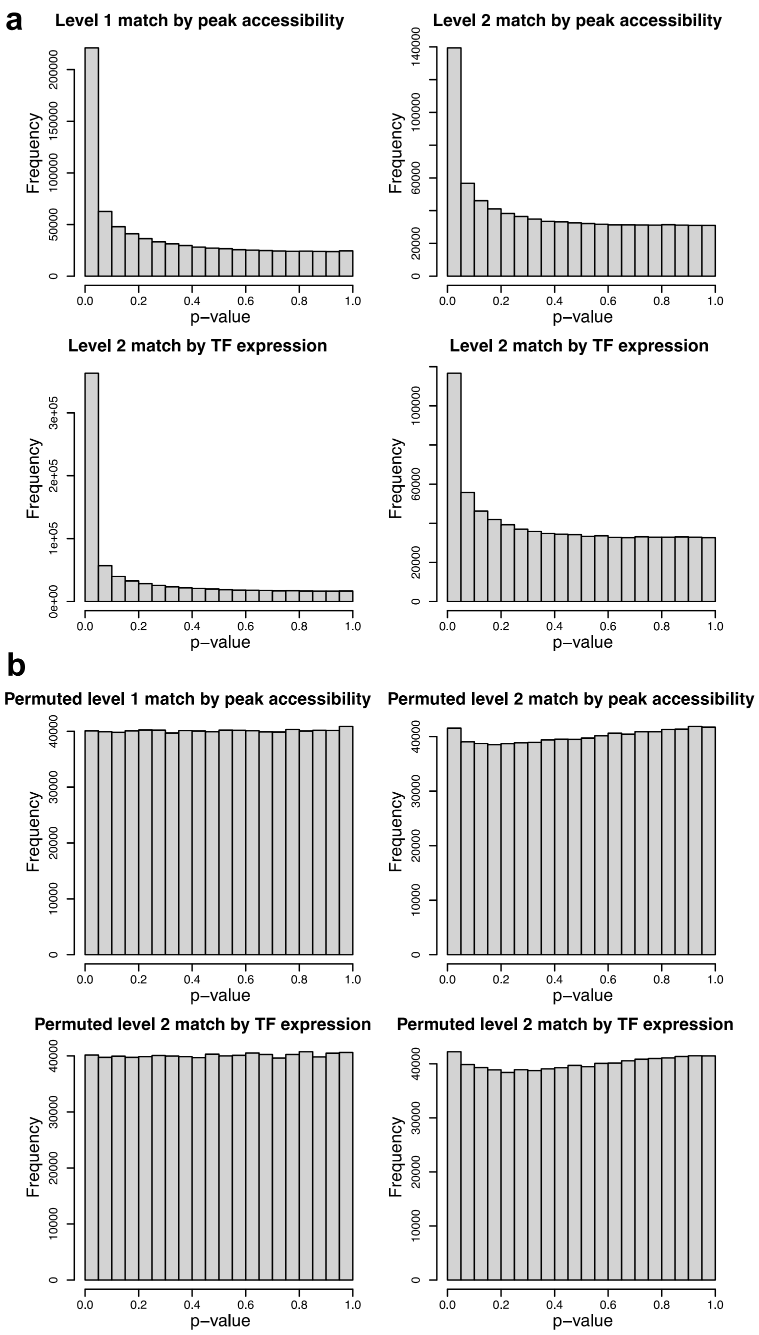

**Supplementary Fig. 13 | Pairwise correlations of genome-wide p-values from different testing schemes.** Transformed $p$-values from marginal association (gene~peak, gene~TF, and peak~TF), linear interaction, TRIPOD's level 1 and level 2 conditional testing matching by peak accessibility and TF expression are shown in the pairwise scatterplots with correlation coefficients shown in the upper triangle. TRIPOD's significant regulatory trios complement both existing approaches based on marginal associations and linear models. Master output is included in Supplementary Table 2.

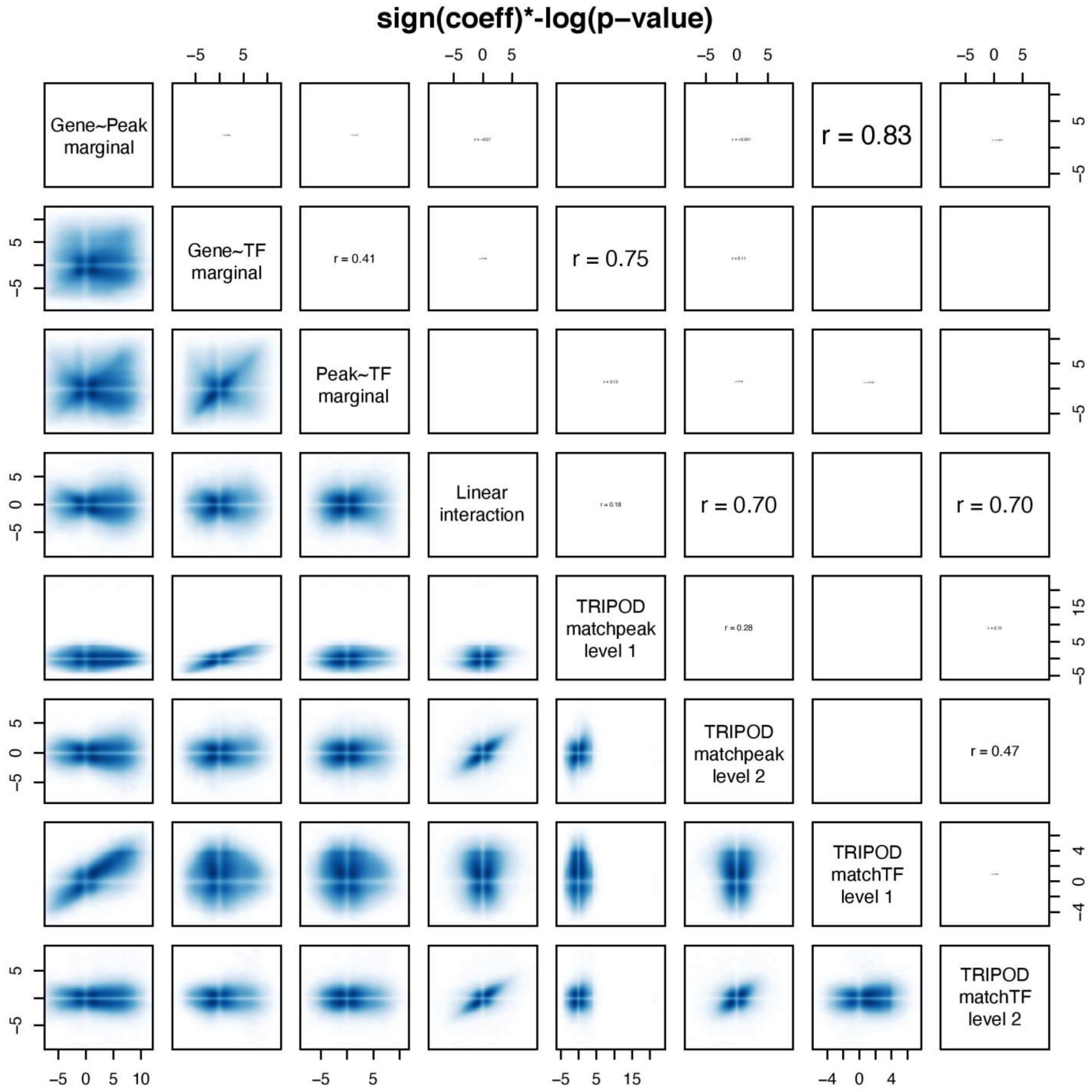

**Supplementary Fig. 14 | TRIPOD's peak-gene validation from TRIPOD's stratified testing scheme.** a, Gene-peak validation using significant trios returned by TRIPOD's level 1 and level 2 testing, matching by peak accessibility and TF expression, respectively. b, Gene-peak validation using significant trios called by TRIPOD but not by random matching (equivalent to random matching). TRIPOD identified additional real regulatory relationships masked by methods based on marginal association testing.

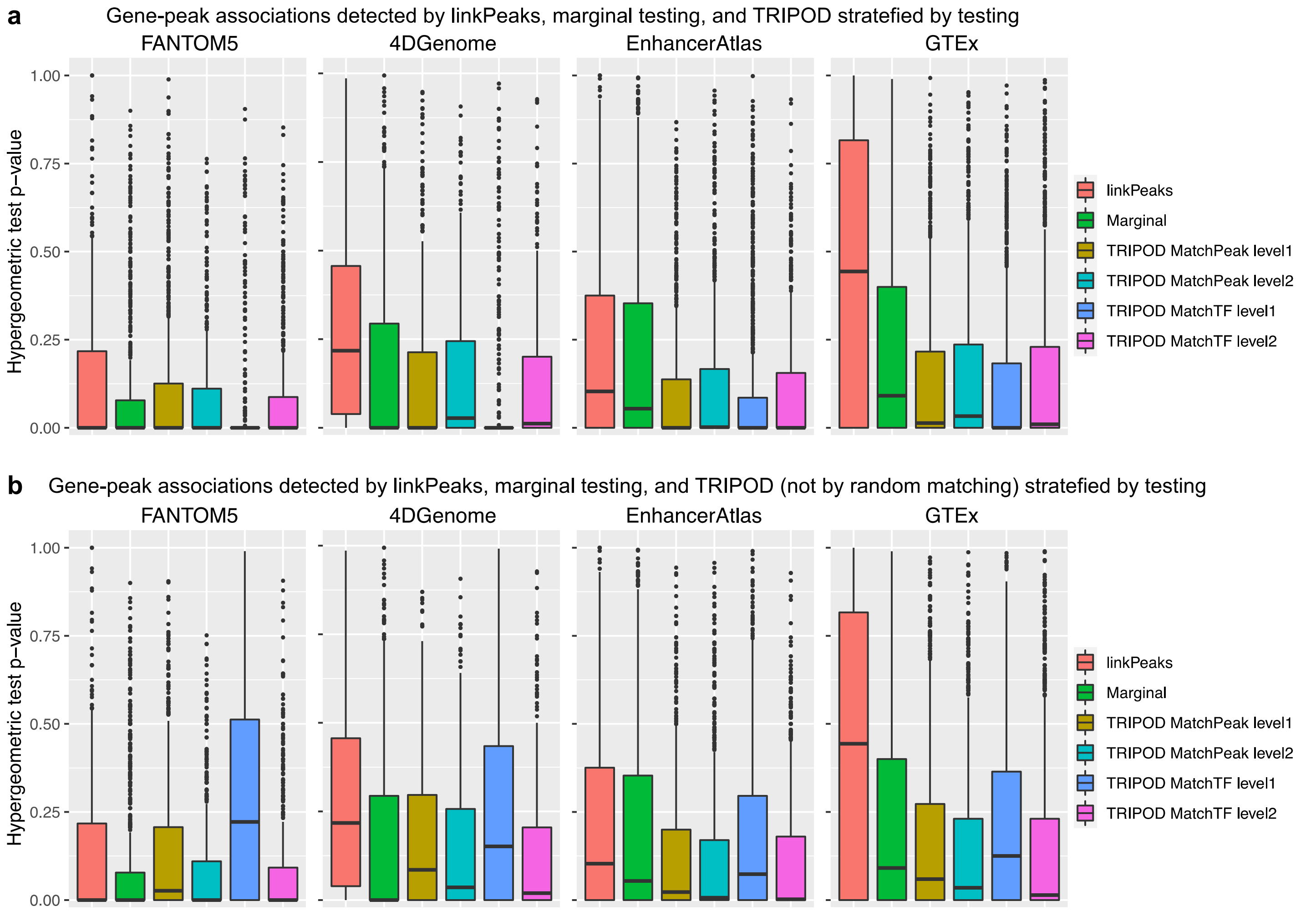

**Supplementary Fig. 15 | Precision and recall rates of TF-gene validation by hTFtarget with different significance levels.** Varying significance threshold alpha is adopted, in addition to the results shown in Fig. 4d.

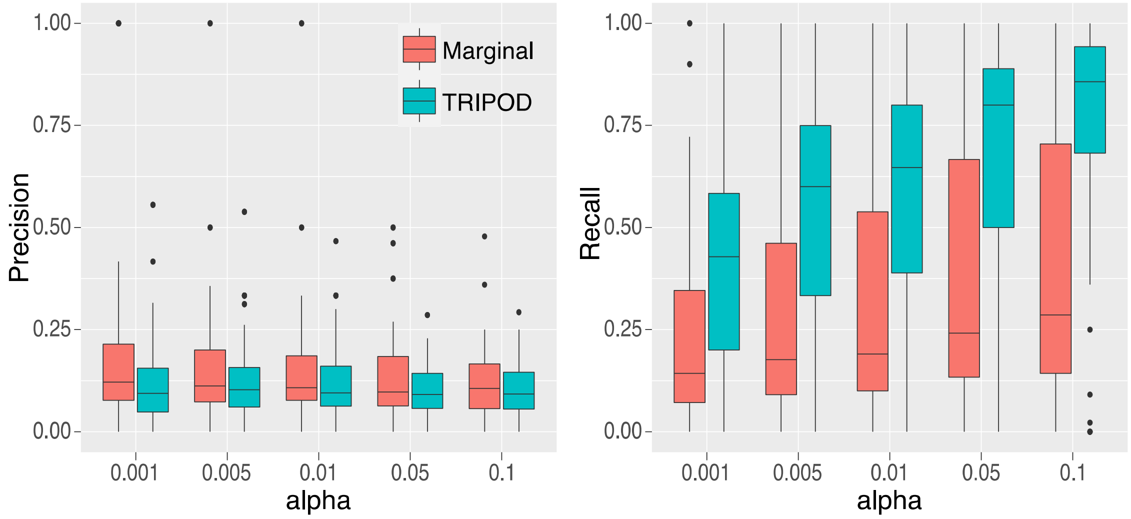

**Supplementary Fig. 16 | Trio regulatory relationships identified by LinkPeaks, marginal association, and TRIPOD, and validation thereof using PLAC-seq data. a**, Bar plots of the numbers of significant regulatory links detected by TRIPOD and marginal associations (FDR < 0.01). The numbers of peak-gene pairs and TF-gene pairs were obtained by collapsing trios by TFs and peaks, respectively. **b**, Heatmap showing the degree of enrichment of ATAC peaks in enhancer-promoter contacts by PLAC-seq[5]. **c**, Venn diagrams of the number of peak-gene pairs captured by PLAC-seq, marginal association between gene expression and peak accessibility, and various models as indicated on the top of each diagram.

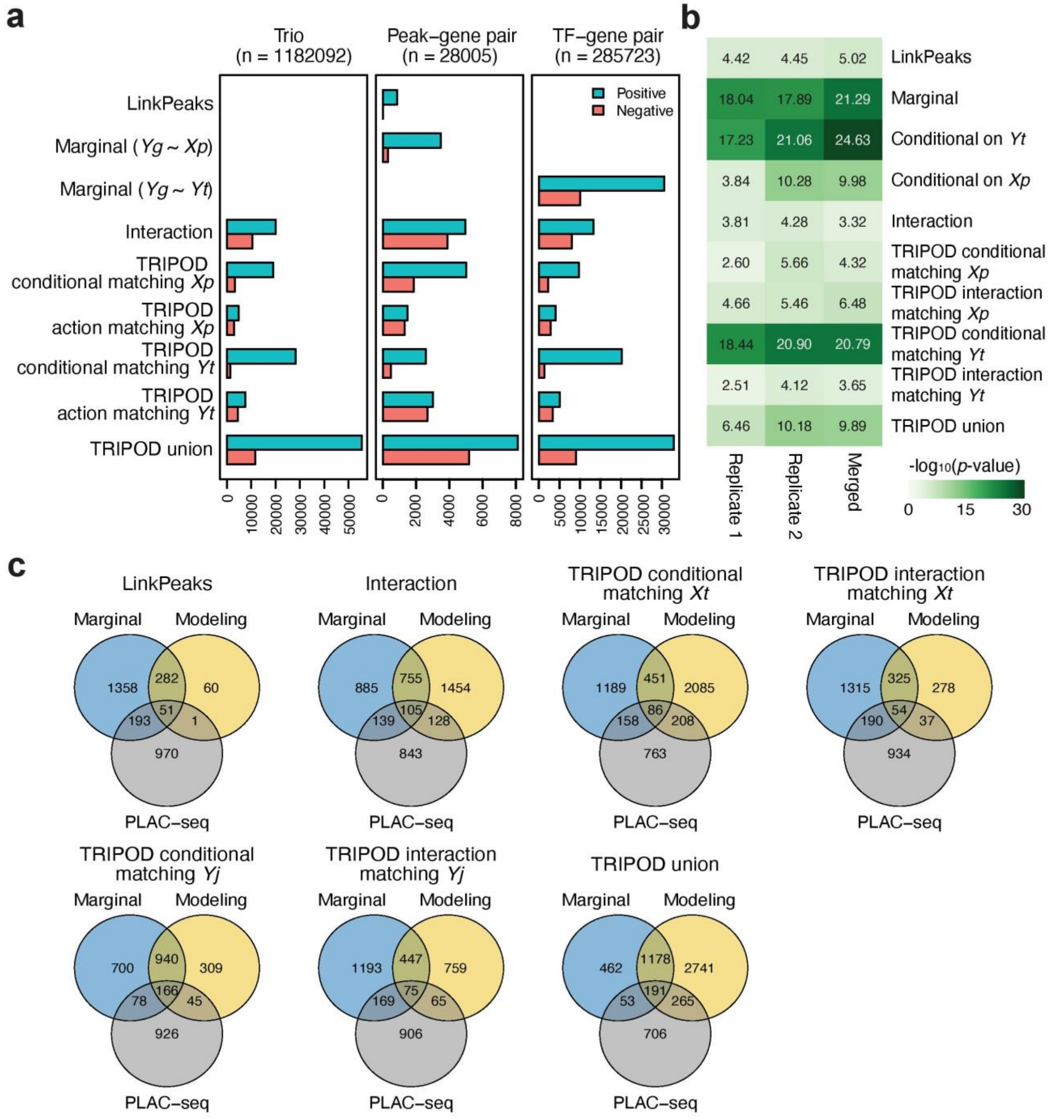

**Supplementary Fig. 17 | Effects of window sizes on the model performance. a**, Heatmap showing the degree of enrichment of ATAC peaks in enhancer-promoter contacts by PLAC-seq[5] with window sizes corresponding to 50, 100, and 200 kb up and downstream from TSS. **b**, Peak-TF validation by ChIP-seq data for Olig2, Nuerog2, Eomes, and Tbr1 (Table 1) at window sizes of 50 kb (**b**), 100 kb (**c**), and 200 kb (**d**). Note that Olig2 ChIP-seq data from mature oligodendrocytes (mOL) serves as a negative control.

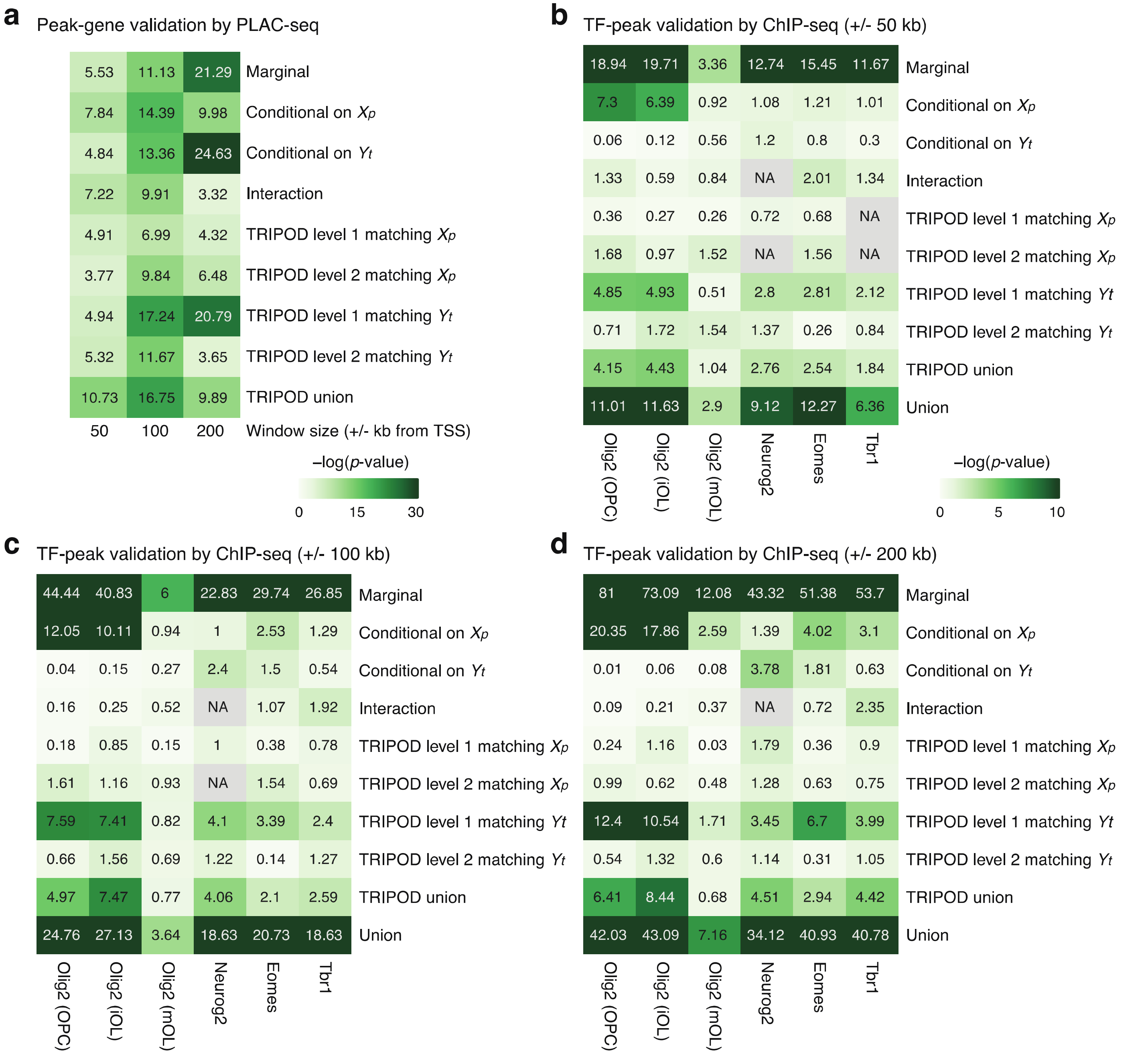

**Supplementary Fig. 18 | Effects of metacell resolutions on the model performance. a**, Heatmap showing the degree of enrichment of ATAC peaks in enhancer-promoter contacts by PLAC-seq[5] with metacells obtained at various clustering resolutions. The window size was set to 200 kb up and downstream of TSS. **b-d**, Peak-TF validation by ChIP-seq data (Table 1) for Olig2, Nuerog2, Eomes, and Tbr1 at clustering resolutions of 10 (**b**), 15 (**c**), and 20 (**d**) with the window size of 200 kb. Note that Olig2 ChIP-seq data from mature oligodendrocytes (mOL) serves as a negative control.

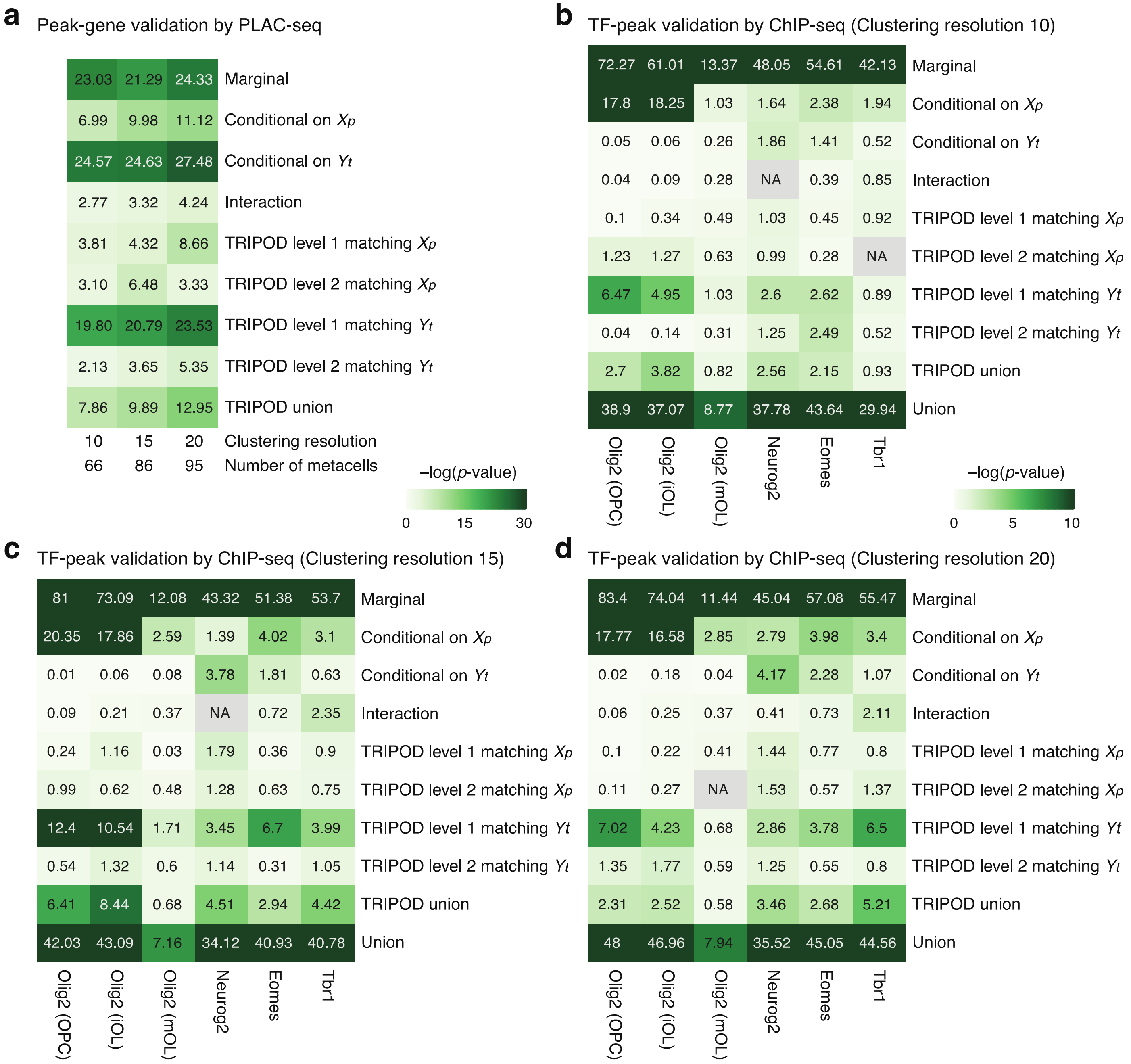

**Supplementary Fig. 19 | Regulatory relationships between neurogenesis and gliogenesis pathways.** The heatmap in the top panel show *p*-values for the marginal associations between TF and gene expression for 11 regulatory relationships indicated in the bottom. Note that the coefficient estimates were positive (i.e., greater than zero) for all cases. The bottom panel shows numbers of trio regulatory relationships with significantly positive coefficients (FDR < 0.01).

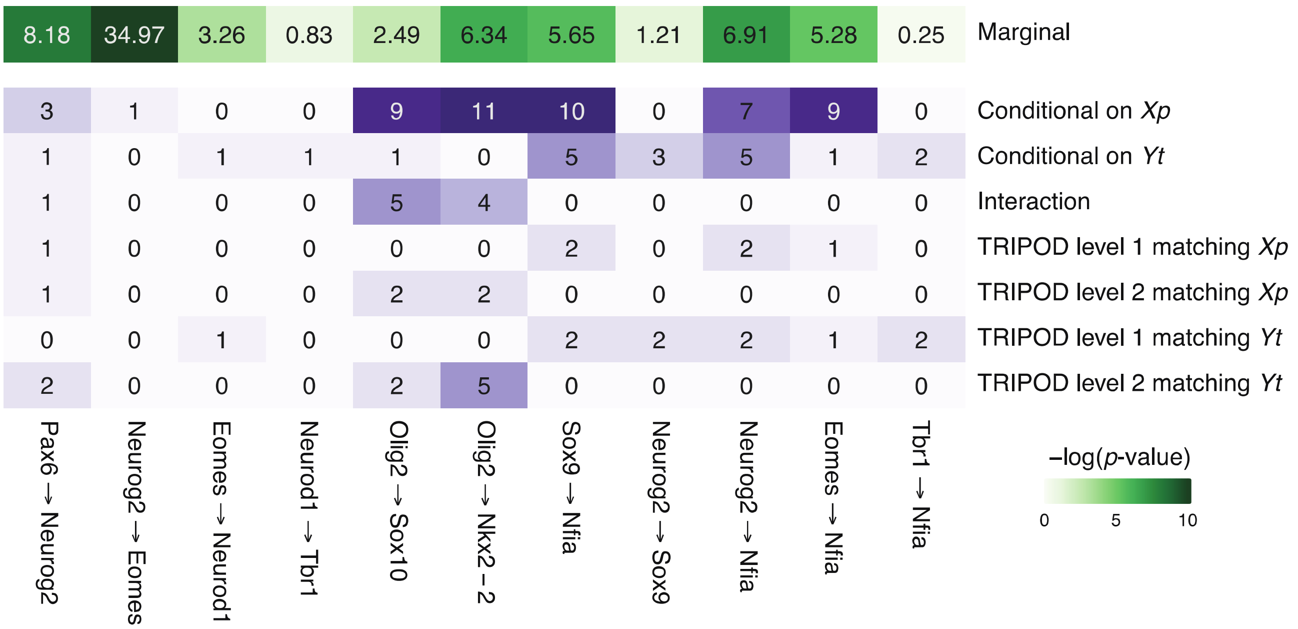

**Supplementary Fig. 20 | Visualization of target gene expression levels, chromatin accessibility in the ATAC peak regions, and TF expression levels for example trios. a-e,** The levels of gene expression (left), peak accessibility (middle), and TF expression (right) are shown on UMAP embeddings for example trios in Fig. 5d.

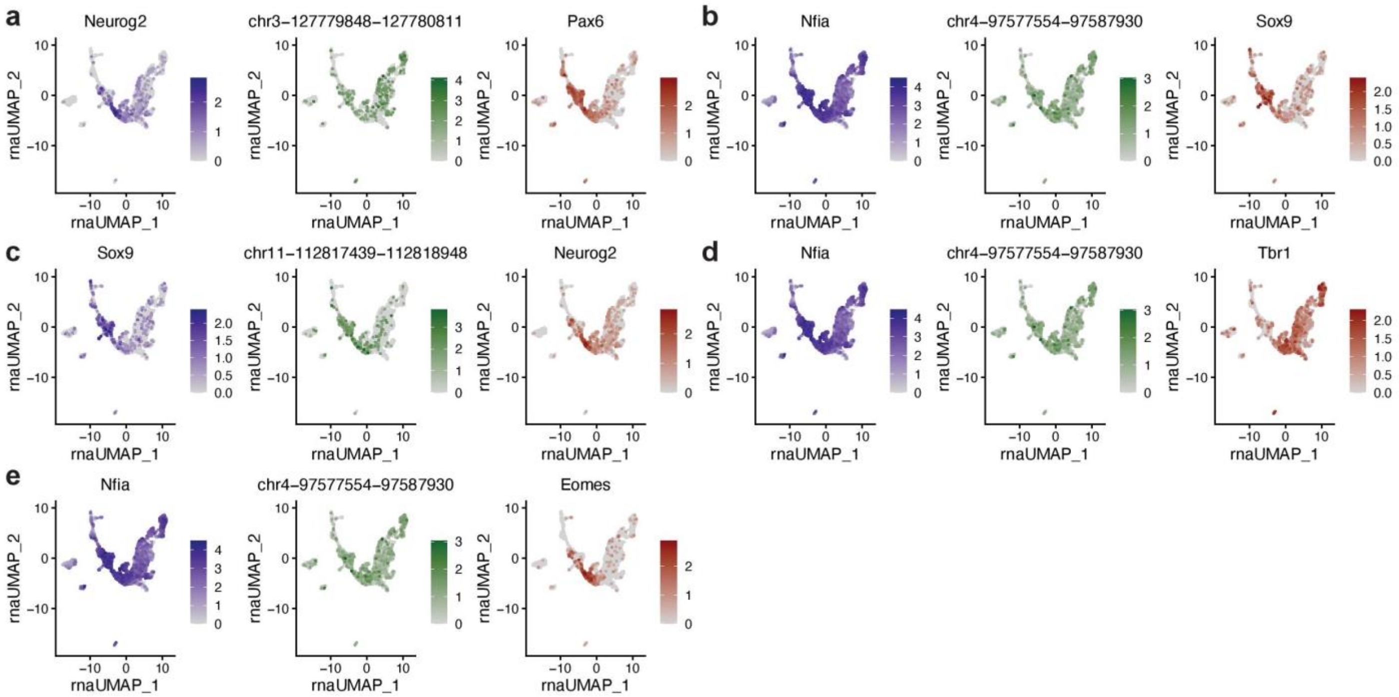

**Supplementary Fig. 21 | Visualization of regulatory links representing possible crosstalks between neurogenesis- and gliogenesis-specific TF cascades. a**, Putative regulation of *Nfia* by Neurog2, Eomes, and Tbr1. **b**, Putative regulation of *Sox9* by Neurog2. ChIP-seq peaks for the indicated TFs are included; arcs represent significant regulatory links inferred by TRIPOD.

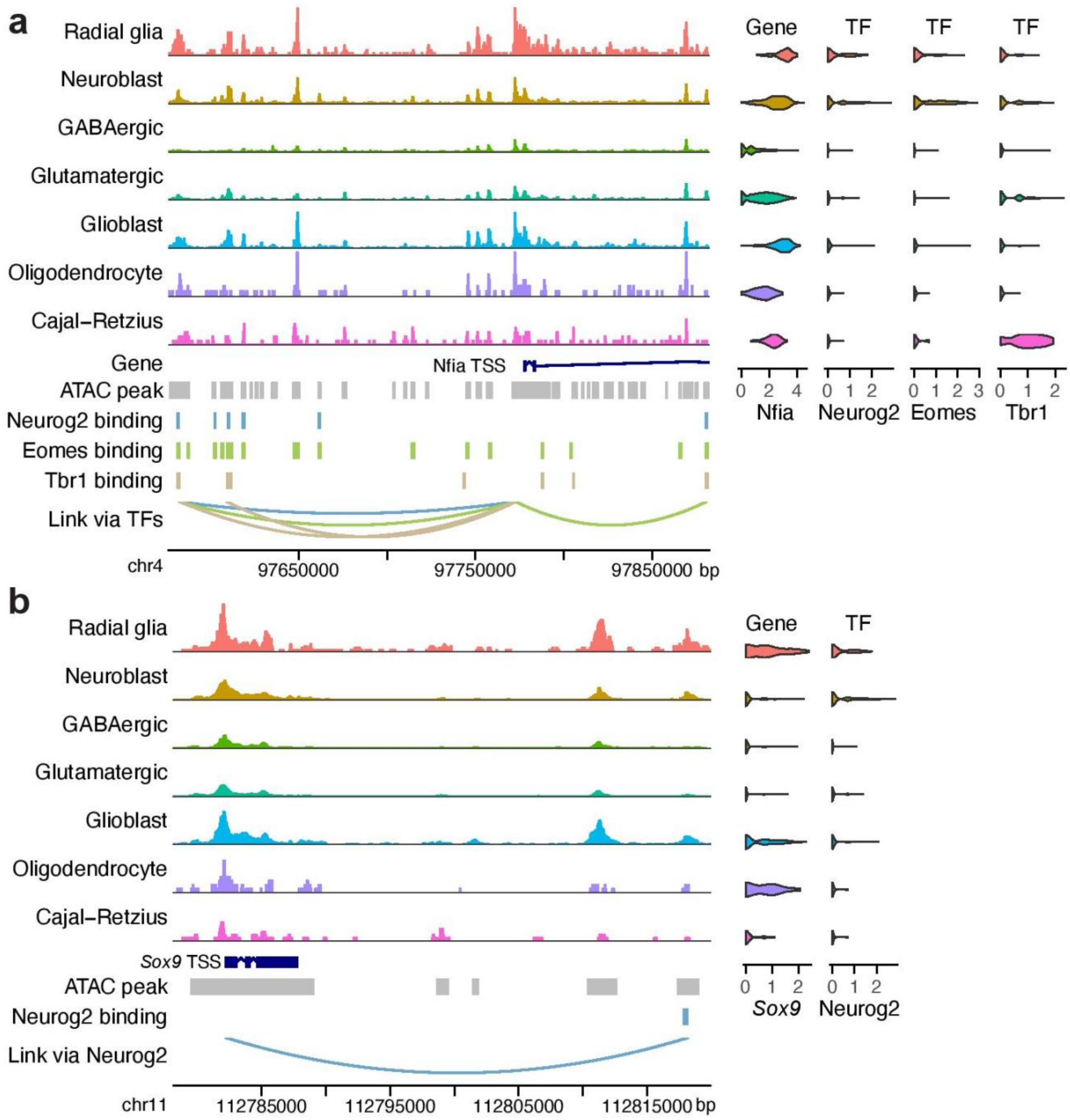

**Supplementary Fig. 22 | Identification of putative cell-type- and cell-state-specific regulation in mouse embryonic brain. a-h**, Visualization of eight example trios from the neurogenesis and gliogenesis TF cascades. The scatter plots (left) show TRIPOD's modeling fitting; the points represent metacells and are colored based on cell types. The middle panels show cell-type-specific $p$-values from the sampling-based influence analyses. The colors on the UMAP embedding (right) correspond to the smoothed $p$-values from the sampling-based influence analyses along the differentiation trajectory. Genomic coordinates for the peaks are from mm10.

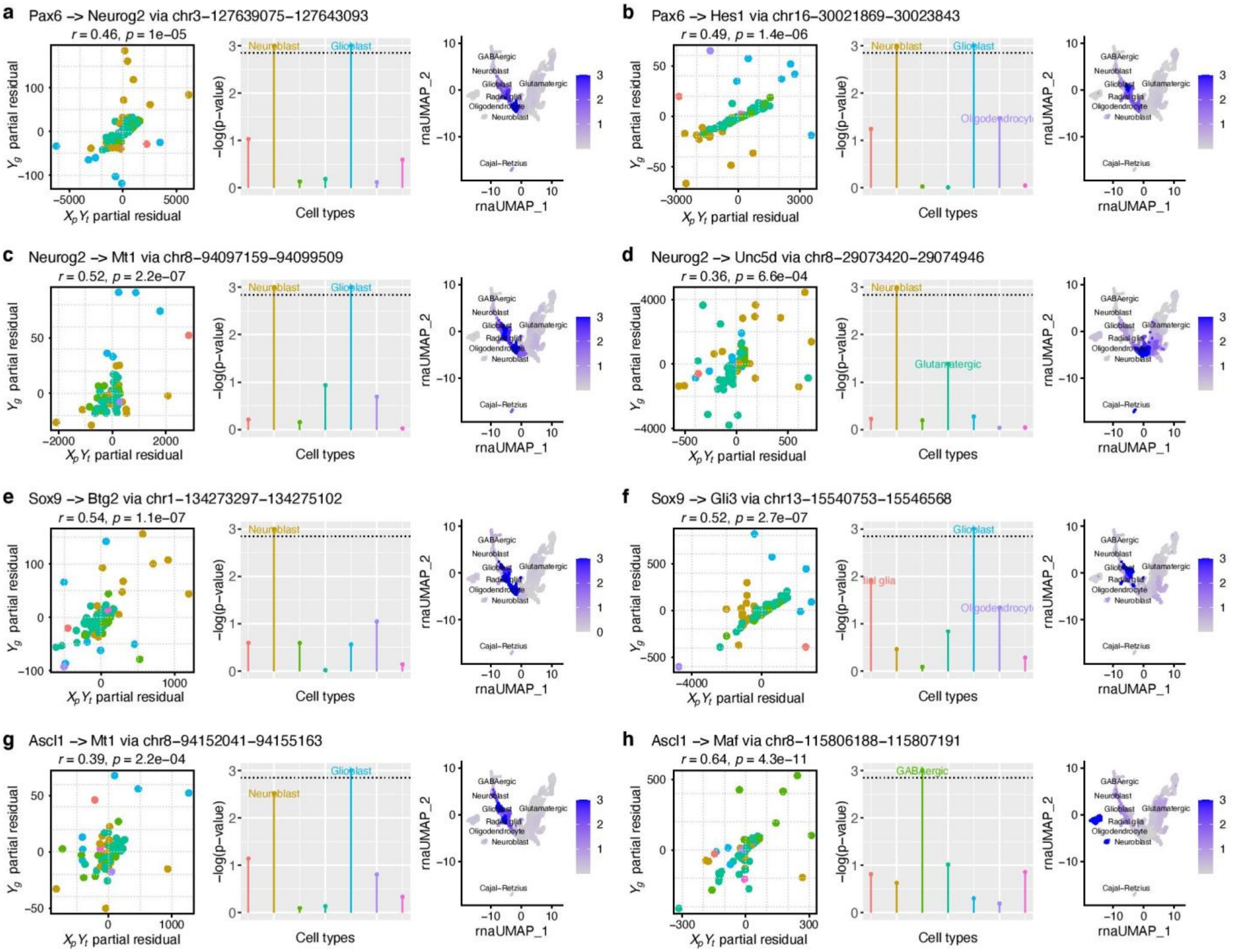

**Supplementary Fig. 23 | Cell-type-specific expression levels of neurogenesis and gliogenesis TFs.** The violin plots show distributions of RNA expression levels of 10 TFs (Pax6, Neurog2, Eomes, Neurod1, Tbr1, Olig2, Sox10, Sox9, Nfia, and Ascl1) that were normalized using sctransform7 and stratified by inferred cell types in the mouse embryonic brain data. The significantly influential cell types identified by TRIPOD are underlined and bolded.

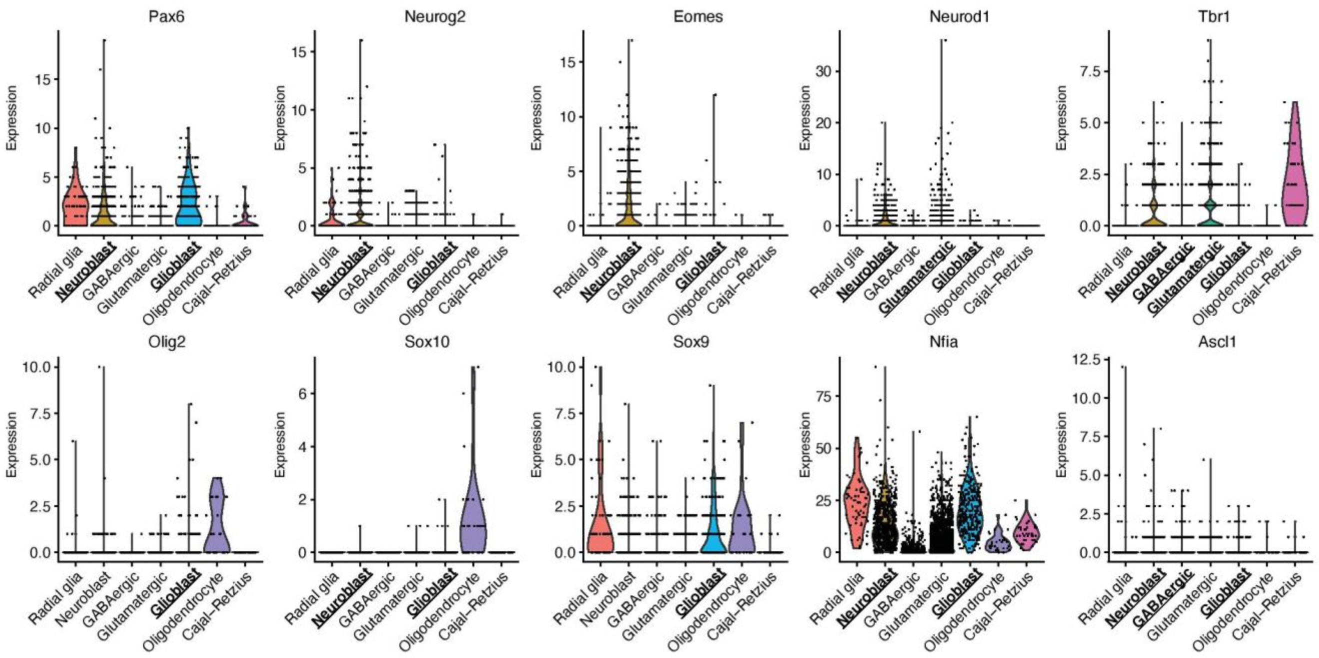

**Supplementary Fig. 24 | Visualization of target gene expression levels, chromatin accessibility in the ATAC peak regions, and TF expression levels for example trios. a-c,** The levels of gene expression (left), peak accessibility (middle), and TF expression (right) are shown on UMAP embeddings for example trios in Fig. 6d,e.

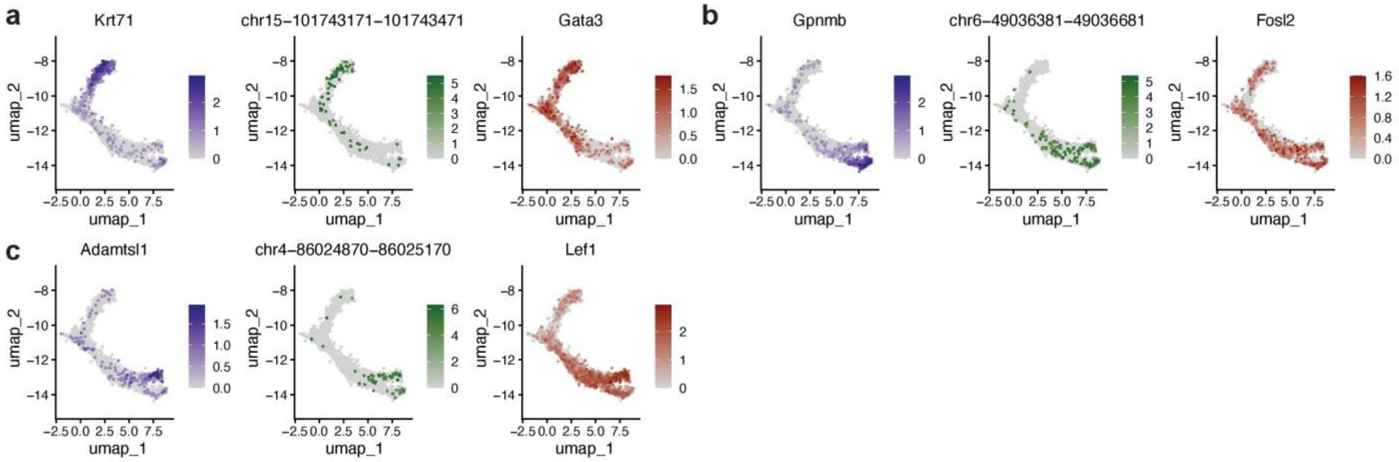

**Supplementary Fig. 25 | Combining accessibilities of ATAC peaks containing the TF binding sites within the window centered at the gene's TSS does not improve EPIC's performance.** In this analysis, we considered pairs of a TF and a target gene, and the pair was called positive if the model fitting yielded a positive coefficient estimate with FDR-adjusted p-value less than 0.01. To evaluate the model performance, we used sets of TF-gene pairs obtained from ChIP-seq of Olig2, Neurog2, Eomes, and Tbr1 (Table 1) as ground truths. Specifically, a true regulatory relationship was assumed if there existed at least one ChIP-seq peak within 200 kb up and downstream of the TSS of the target gene. The resultant receiver operator curves (ROCs) are in red. For comparison, ROCs for the models where individual peak regions were analyzed separately are also shown in blue. In this case, lists of significant regulatory trios (FDR < 0.01) were collapsed to obtain lists of TF-gene pairs.

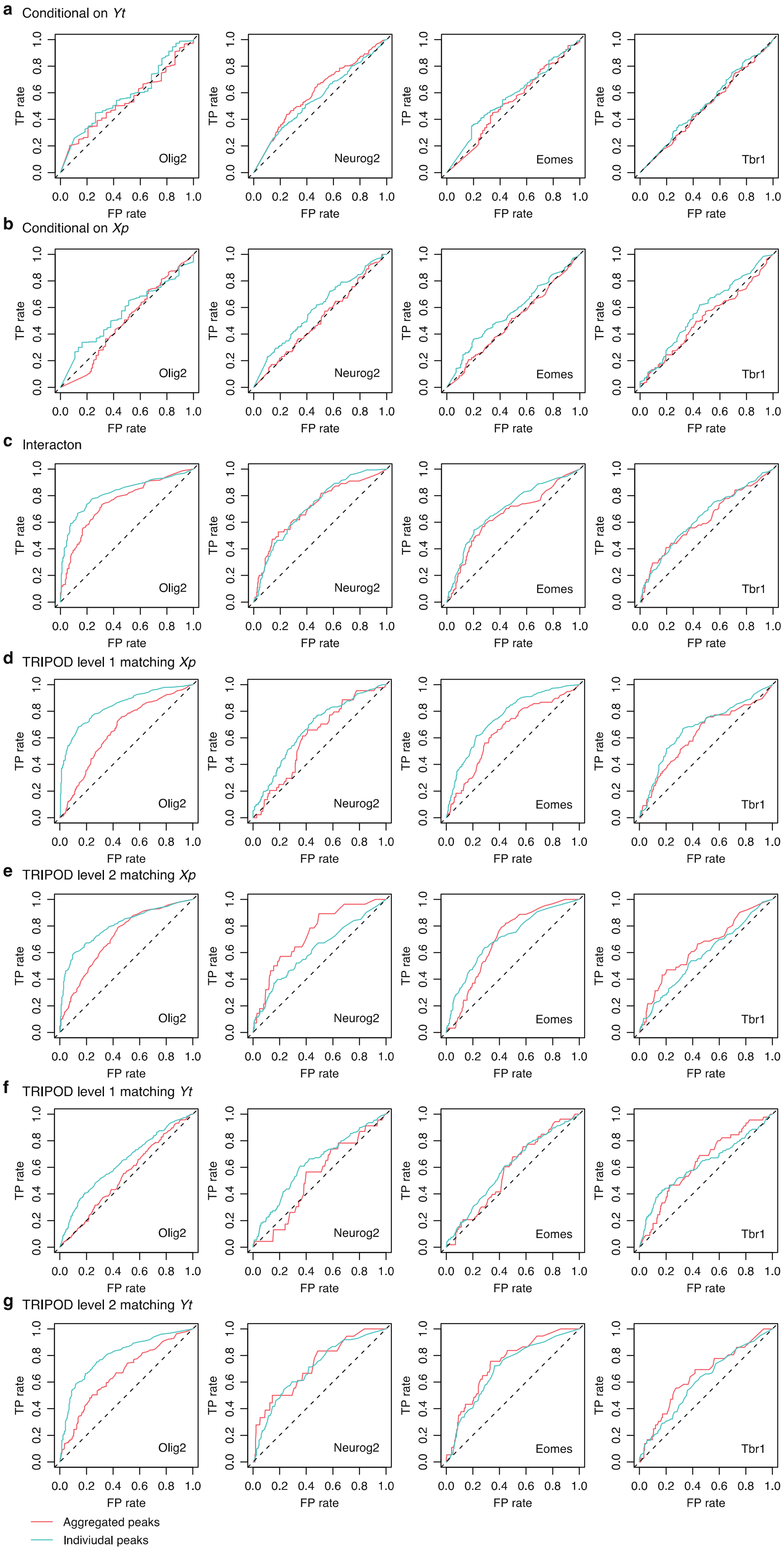

a Conditional on *Yt*
b Conditional on *Xp*
c Interacton
d TRIPOD level 1 matching *Xp*
e TRIPOD level 2 matching *Xp*
f TRIPOD level 1 matching *Yt*
g TRIPOD level 2 matching *Yt*
Olig2
Neurog2
Eomes
Tbr1
TP rate
FP rate
0.0 0.2 0.4 0.6 0.8 1.0
Aggregated peaks
Indiviudal peaks

**Supplementary Fig. 26 | Assessing linear models for detecting conditional associations**. Scatter plots comparing fittings from linear models and TRIPOD's level 1 and level 2 testing for example trios in **a**, PBMC and **b**, mouse skin. Genomic coordinates for the peaks are from hg38 for human and mm10 for mouse.

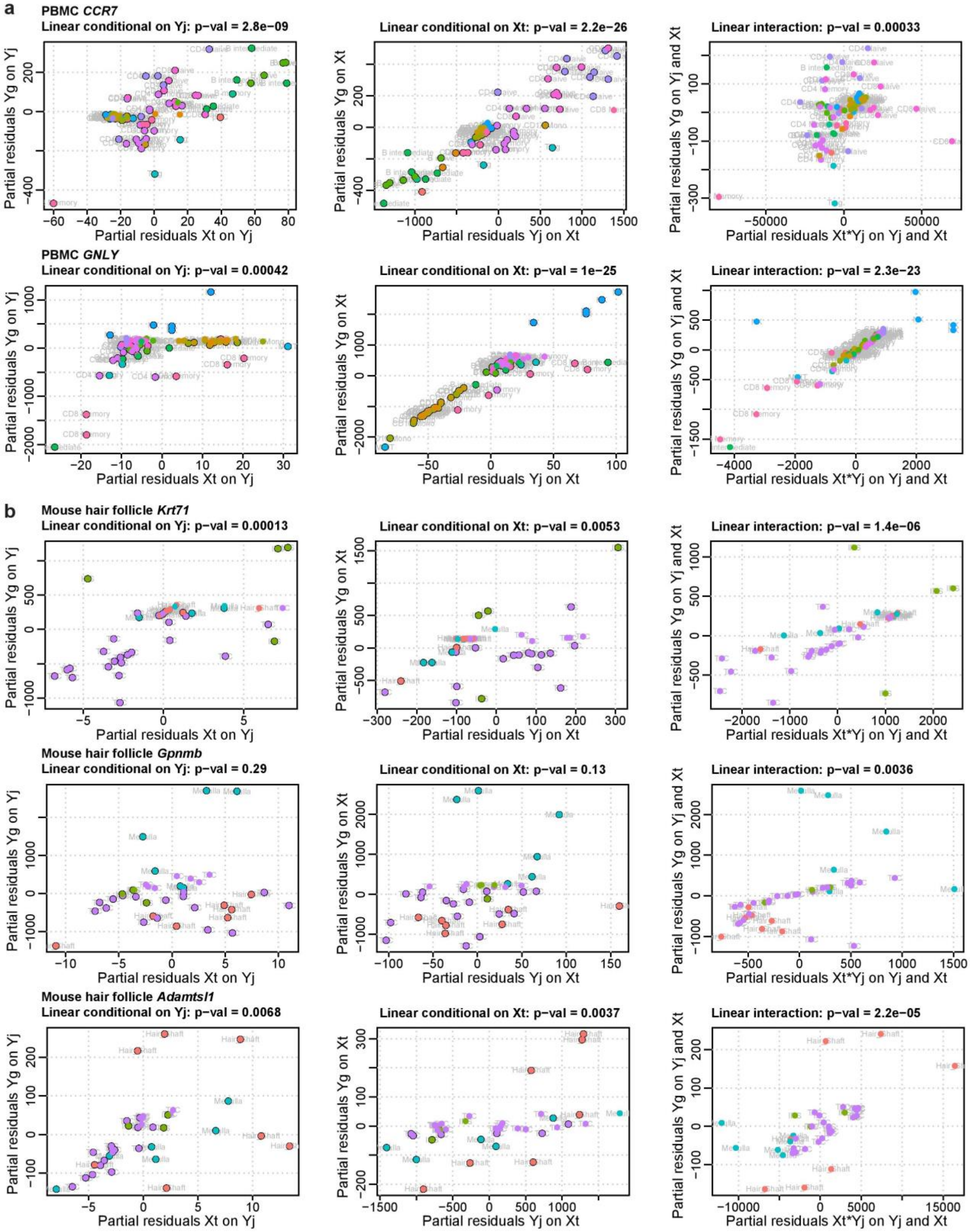

**Supplementary Fig. 27 | Comparison of estimated coefficients from linear model and TRIPOD'S nonparametric model. a-d**, Pairwise scatter plots comparing transformed coefficients from linear model and TRIPOD's nonparametric model for representative target genes, *CCR7*, *GNLY*, *FCGR3A*, and *MS4A1* from the PBMC data. $\gamma$ denotes the coefficient for the interaction between TF expression and peak accessibility; $\alpha$ and $\beta$ denote the coefficients for the partial gene-peak and gene-TF correlations, respectively. The three coefficients are fit using the linear model and TRIPOD's level 1 and level 2 testing, and their estimates are correlated on the global scale. However, the actual call sets are different, and the underlying models and assumptions are different.

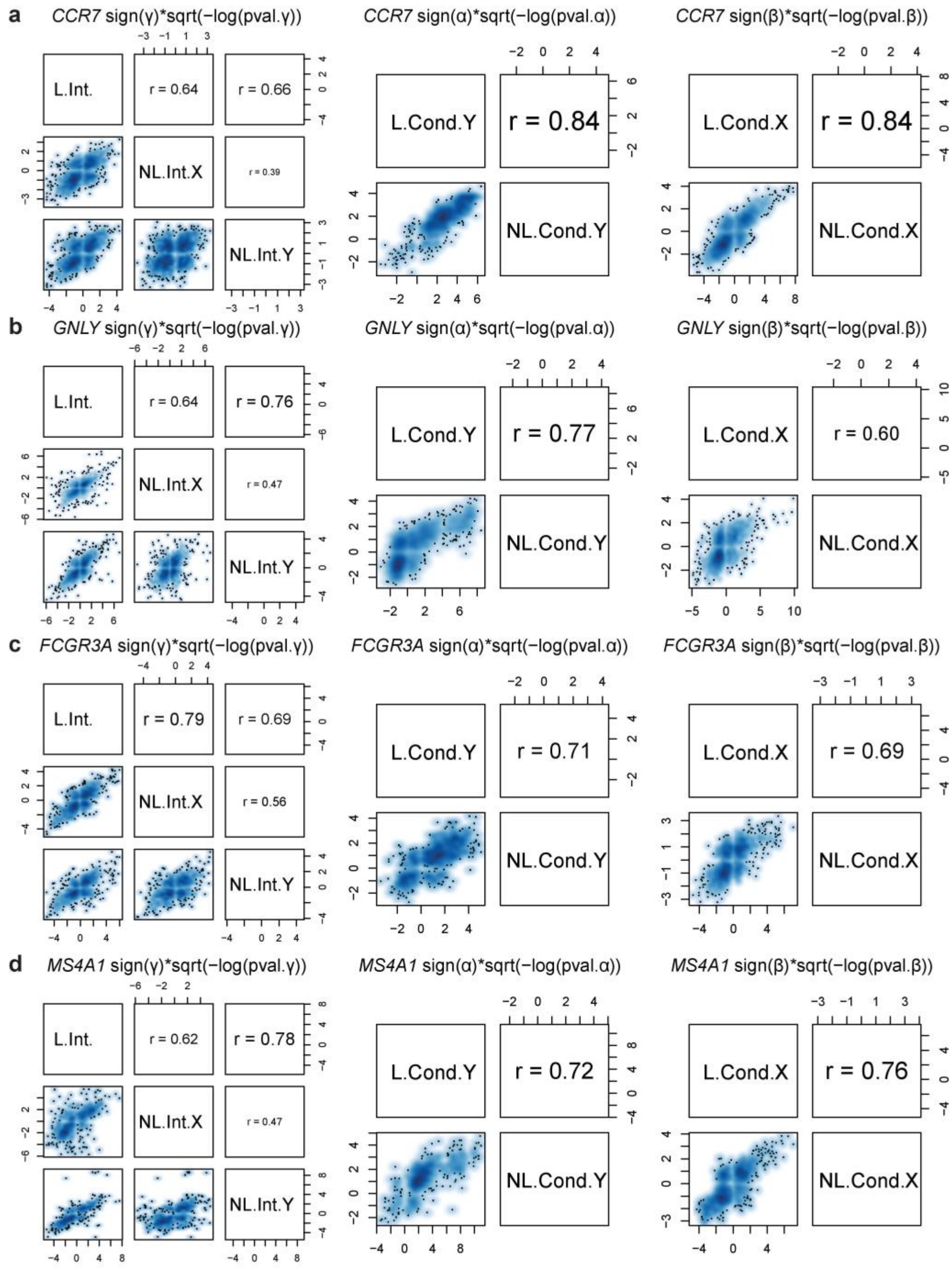

**Supplementary Fig. 28 | Comparison of the trio regulatory relationships identified by linear model and TRIPOD's nonparametric model. a-d**, Venn diagrams of the number of significant trios detected by linear and TRIPOD models for representative target genes, *CCR7*, *GNLY*, *FCGR3A*, and *MS4A1*. The left, middle, and right panels contain results from the interaction models, the models conditional on TF expression $Y_t$, and the models conditional on peak accessibility $X_p$. L.int., TRIPOD.Int.Y, TRIPOD.Int.X, L.Cond.Y, TRIPOD.Cond.Y, L.Cond.X, and TRIPOD.Cond.X represent linear interaction model, nonparametric interaction model matching by TF expression (TRIPOD level 2 test), nonparametric interaction model matching by peak accessibility (TRIPOD level 2 test), linear model conditional on TF expression, nonparametric conditional model matching by TF expression (TRIPOD level 1 test), linear model conditional on peak accessibility, and nonparametric conditional model matching by peak accessibility (TRIPOD level 1 test), respectively.

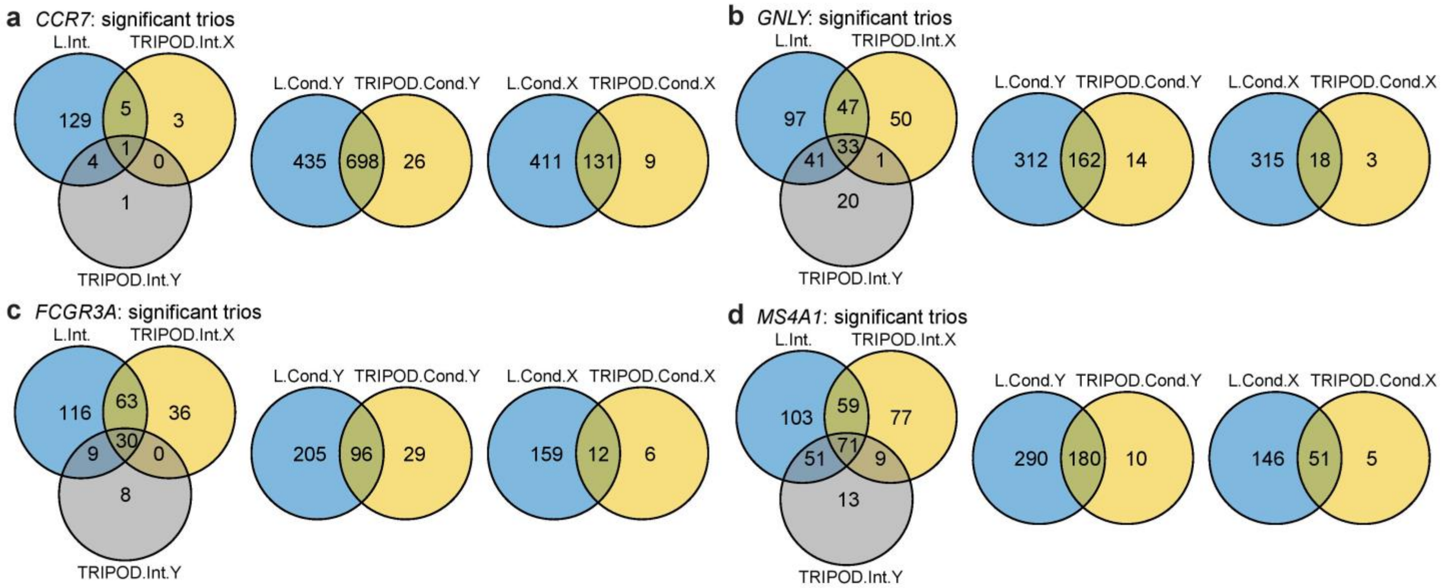

### Supplementary Tables

**Supplementary Table 1 | Single-cell RNA and ATAC multiomic datasets.** Data adopted in this study from different tissues, organisms, and cell types using different protocols are summarized. The number of cells, peaks, and genes, as well as RNA read counts and ATAC read counts **a**, before and **b**, after quality control (QC) are summarized. Sources of data are also provided.

(A)

| Organism | Tissue | Protocol | PMID or source | Before QC | | | | |
|---|---|---|---|---|---|---|---|---|
| | | | | Number of cells | Number of peaks | Number of genes | Mean cov. ATAC | Mean cov. RNA |
| *Homo sapiens* | PBMC | 10X Genomics Multiome | 10X Genomics | 11909 | 108377 | 36601 | 20434 | 4402 |
| *Homo sapiens* | PBMC | 10X Genomics Multiome | 10X Genomics | 2714 | 156607 | 36601 | 20661 | 4344 |
| *Mus musculus* | Embryonic brain at day 18 | 10X Genomics Multiome | 10X Genomics | 4881 | 139083 | 32285 | 20989 | 9486 |
| *Mus musculus* | Skin late anagen | SHARE-seq | 33098772 | 34774 | 344592 | 23296 | 4231 | 1259 |
| *Mus musculus* | Adult brain cerebral cortex | SNARE-seq | 31611697 | 10309 | 244544 | 33160 | 2656 | 1549 |
| *Mus musculus* | Adult brain cerebral cortex | PAIRED-seq | 31695190 | 15191 | 2614863 | 29624 | 1600 | 459 |

(B)

| Organism | Tissue | Protocol | PMID or source | After QC | | | | |
|---|---|---|---|---|---|---|---|---|
| | | | | Number of cells | Number of peaks | Number of genes | Mean cov. ATAC | Mean cov. RNA |
| *Homo sapiens* | PBMC | 10X Genomics Multiome | 10X Genomics | 7790 | 103755 | 14508 | 9611 | 3352 |
| *Homo sapiens* | PBMC | 10X Genomics Multiome | 10X Genomics | 2206 | 106056 | 36601 | 13534 | 4425 |
| *Mus musculus* | Embryonic brain at day 18 | 10X Genomics Multiome | 10X Genomics | 3962 | 139083 | 14476 | 12520 | 8293 |
| *Mus musculus* | Skin late anagen | SHARE-seq | 33098772 | 29308 | 343783 | 15086 | 17886 | 1080 |
| *Mus musculus* | Adult brain cerebral cortex | SNARE-seq | 31611697 | 7533 | 244544 | 15332 | 12534 | 2053 |
| *Mus musculus* | Adult brain cerebral cortex | PAIRED-seq | 31695190 | 11292 | 118337 | 17858 | 3014 | 387 |

**Supplementary Table 2 | Master output from PBMC multiomic analysis.** Testing results from marginal association testing, linear conditional and interaction model, as well as TRIPOD's two levels of testing are returned. Separately attached as an excel file.

**Supplementary Table 3 | Precision, recall, and joint F1 score with varying significance levels for TF-gene validation by knockTF.** Varying significance level alpha is adopted, in addition to the fixed alpha shown in Fig. 4c. TRIPOD outperforms methods based on marginal association testing.

| TF (knockTF) | alpha | Precision | | Recall | | F1 score | |
|---|---|---|---|---|---|---|---|
| | | Marginal | TRIPOD | Marginal | TRIPOD | Marginal | TRIPOD |
| GATA3 | 0.001 | 0.0059 | 0.0108 | 0.0024 | 0.0049 | 0.0035 | 0.0067 |
| | 0.005 | 0.0051 | 0.0099 | 0.0024 | 0.0073 | 0.0033 | 0.0084 |
| | 0.01 | 0.005 | 0.0111 | 0.0024 | 0.0098 | 0.0033 | 0.0104 |
| | 0.05 | 0.0086 | 0.0097 | 0.0049 | 0.0122 | 0.0062 | 0.0108 |
| JUN | 0.001 | 0.1256 | 0.1445 | 0.0351 | 0.0437 | 0.0549 | 0.0671 |
| | 0.005 | 0.1244 | 0.123 | 0.0366 | 0.0487 | 0.0565 | 0.0698 |
| | 0.01 | 0.1238 | 0.1156 | 0.0373 | 0.0495 | 0.0573 | 0.0693 |
| | 0.05 | 0.1219 | 0.116 | 0.0387 | 0.0609 | 0.0588 | 0.0799 |
| MAF | 0.001 | 0 | 0.0917 | 0 | 0.0267 | 0 | 0.0414 |
| | 0.005 | 0 | 0.0906 | 0 | 0.0352 | 0 | 0.0507 |
| | 0.01 | 0 | 0.0871 | 0 | 0.0377 | 0 | 0.0526 |
| | 0.05 | 0.0071 | 0.0774 | 0.0012 | 0.0437 | 0.0021 | 0.0559 |
| MYB | 0.001 | 0 | 0.1468 | 0 | 0.0424 | 0 | 0.0658 |
| | 0.005 | 0.037 | 0.1324 | 0.0027 | 0.0716 | 0.005 | 0.0929 |
| | 0.01 | 0.0256 | 0.1265 | 0.0027 | 0.0849 | 0.0048 | 0.1016 |
| | 0.05 | 0.0282 | 0.1275 | 0.0053 | 0.1353 | 0.0089 | 0.1313 |
| NFATC3 | 0.001 | 0.0403 | 0.0824 | 0.0047 | 0.0339 | 0.0084 | 0.0481 |
| | 0.005 | 0.0357 | 0.0835 | 0.0057 | 0.0452 | 0.0098 | 0.0587 |
| | 0.01 | 0.0291 | 0.0821 | 0.0057 | 0.049 | 0.0095 | 0.0614 |
| | 0.05 | 0.0247 | 0.0786 | 0.0066 | 0.0594 | 0.0104 | 0.0676 |
| NFKB1 | 0.001 | 0 | 0.0076 | 0 | 0.0105 | 0 | 0.0088 |
| | 0.005 | 0 | 0.006 | 0 | 0.0105 | 0 | 0.0076 |
| | 0.01 | 0 | 0.0072 | 0 | 0.014 | 0 | 0.0095 |
| | 0.05 | 0 | 0.0087 | 0 | 0.021 | 0 | 0.0123 |
| ZNF148 | 0.001 | 0 | 0.0013 | 0 | 0.027 | 0 | 0.0025 |
| | 0.005 | 0 | 0.0035 | 0 | 0.0811 | 0 | 0.0068 |
| | 0.01 | 0 | 0.0034 | 0 | 0.0811 | 0 | 0.0065 |
| | 0.05 | 0 | 0.003 | 0 | 0.0811 | 0 | 0.0058 |

**Supplementary Table 4 | ChIP-seq data of human blood (B lymphocyte, T lymphocyte, and monocyte).** Non-cancerous and cell-type-specific ChIP-seq data of human blood were downloaded from the Cistrome[8] portal to validate the peak-TF links in the PBMC single-cell multiomic data. PCR bottleneck coefficient (PBC) is used to estimate the rate of read duplication through PCR amplification; a good PBC score is ≥ 80%. PeaksFoldChangeAbove10 contains the number of peaks called by MACS2[9] with a fold change above 10. FRiP is used for evaluating the signal-to-noise ratio; a good FRiP score is ≥ 1%. PeaksUnionDHSRatio is the percentage of the merged top 5000 peaks (ordered by MACS2 $q$-value) that overlap with union DHS regions; this is expected to be ≥ 70%.

| DCid | Species | GSMID | Factor | Cell_line | Cell_type | Tissue_type | FastQC | UniquelyMappedRatio | PBC | PeaksFoldChangeAbove10 | FRiP | PeaksUnionDHSRatio |
|---|---|---|---|---|---|---|---|---|---|---|---|---|
| 40022 | Homo sapiens | GSM1121094 | BRD4 | None | B Lymphocyte | Blood | 38 | 0.8193 | 0.955 | 1615 | 0.1565535 | 0.9436 |
| 40053 | Homo sapiens | GSM1121098 | CDK7 | None | B Lymphocyte | Blood | 38 | 0.7267 | 0.962 | 1230 | 0.05939275 | 0.9608 |
| 40226 | Homo sapiens | GSM1195559 | CDK9 | None | B Lymphocyte | Blood | 38 | 0.7745 | 0.937 | 4517 | 0.09261 | 0.9696 |
| 45178 | Homo sapiens | GSM1003474 | CTCF | None | B Lymphocyte | Blood | 38 | 0.7908 | 0.904 | 27265 | 0.305596 | 0.9758 |
| 45179 | Homo sapiens | GSM1003476 | H2AZ | None | B Lymphocyte | Blood | 37 | 0.7298 | 0.94 | 2524 | 0.1956965 | 0.975 |
| 40215 | Homo sapiens | GSM1195560 | IRF4 | None | B Lymphocyte | Blood | 38 | 0.8383 | 0.948 | 8535 | 0.13387325 | 0.9422 |
| 40216 | Homo sapiens | GSM1195557 | MED1 | None | B Lymphocyte | Blood | 38 | 0.5248 | 0.844 | 4058 | 0.14310625 | 0.9664 |
| 40233 | Homo sapiens | GSM1195555 | MED1 | None | B Lymphocyte | Blood | 38 | 0.6613 | 0.911 | 9273 | 0.2180545 | 0.954 |
| 40234 | Homo sapiens | GSM1195556 | MED1 | None | B Lymphocyte | Blood | 38 | 0.8217 | 0.963 | 4479 | 0.1278495 | 0.9472 |
| 5967 | Homo sapiens | GSM762709 | MYC | None | B Lymphocyte | Blood | 29 | 0.7391 | 0.988 | 1211 | 0.048873002 | 0.9576 |
| 33439 | Homo sapiens | GSM971344 | POLR2A | None | B Lymphocyte | Blood | 30 | 0.7135 | 0.922 | 4948 | 0.1402875 | 0.9746 |
| 33434 | Homo sapiens | GSM971343 | SMARCA4 | None | B Lymphocyte | Blood | 30 | 0.6431 | 0.909 | 2353 | 0.0451365 | 0.964 |
| 45444 | Homo sapiens | GSM1003508 | CTCF | None | Monocyte | Blood | 37 | 0.7086 | 0.948 | 23384 | 0.243658 | 0.9718 |
| 45441 | Homo sapiens | GSM1003548 | H2AZ | None | Monocyte | Blood | 37 | 0.8274 | 0.898 | 1674 | 0.11932275 | 0.975 |
| 41301 | Homo sapiens | GSM1057025 | IRF1 | None | Monocyte | Blood | 38 | 0.8155 | 0.932 | 11224 | 0.07846625 | 0.9264 |
| 41302 | Homo sapiens | GSM1057026 | IRF1 | None | Monocyte | Blood | 38 | 0.8212 | 0.943 | 3018 | 0.028401 | 0.945 |
| 41303 | Homo sapiens | GSM1057027 | IRF1 | None | Monocyte | Blood | 38 | 0.8198 | 0.877 | 5920 | 0.04798775 | 0.9316 |
| 81223 | Homo sapiens | GSM2687534 | RUNX1 | None | Monocyte | Blood | 38 | 0.7012 | 0.919 | 9496 | 0.31398225 | 0.9418 |
| 85986 | Homo sapiens | GSM2804465 | SPI1 | None | Monocyte | Blood | 39 | 0.7064 | 0.944 | 1283 | 0.038436584 | 0.9426 |
| 41287 | Homo sapiens | GSM1057011 | STAT1 | None | Monocyte | Blood | 39 | 0.8088 | 0.993 | 6006 | 0.11584575 | 0.976 |
| 41288 | Homo sapiens | GSM1057012 | STAT1 | None | Monocyte | Blood | 39 | 0.7591 | 0.995 | 1501 | 0.0587395 | 0.9728 |
| 41289 | Homo sapiens | GSM1057013 | STAT1 | None | Monocyte | Blood | 39 | 0.8118 | 0.993 | 11395 | 0.15686975 | 0.9774 |
| 82662 | Homo sapiens | GSM2679938 | T | None | Monocyte | Blood | 39 | 0.8392 | 0.992 | 11428 | 0.14325775 | 0.9586 |
| 81224 | Homo sapiens | GSM2687535 | TET2 | None | Monocyte | Blood | 38 | 0.6452 | 0.79 | 2851 | 0.22890725 | 0.7862 |
| 36301 | Homo sapiens | GSM823379 | BRD4 | None | T Lymphocyte | Blood | 39 | 0.6184 | 0.803 | 1025 | 0.02953575 | 0.9652 |
| 38389 | Homo sapiens | GSM1022944 | BRD4 | None | T Lymphocyte | Blood | 38 | 0.7591 | 0.968 | 1092 | 0.04322775 | 0.9644 |
| 3060 | Homo sapiens | GSM325895 | CTCF | None | T Lymphocyte | Blood | 29 | 0.5214 | 0.902 | 15633 | 0.253560996 | 0.9716 |
| 44090 | Homo sapiens | GSM1056928 | ETS1 | None | T Lymphocyte | Blood | 37 | 0.5843 | 0.965 | 1319 | 0.0132305 | 0.96 |
| 44092 | Homo sapiens | GSM1056930 | ETS1 | None | T Lymphocyte | Blood | 39 | 0.803 | 0.909 | 3052 | 0.0402015 | 0.9594 |
| 44093 | Homo sapiens | GSM1056931 | ETS1 | None | T Lymphocyte | Blood | 38 | 0.7048 | 0.905 | 1559 | 0.02145925 | 0.956 |
| 44094 | Homo sapiens | GSM1056932 | ETS1 | None | T Lymphocyte | Blood | 39 | 0.7582 | 0.903 | 2587 | 0.039202 | 0.9608 |
| 4459 | Homo sapiens | GSM393968 | POLR2A | None | T Lymphocyte | Blood | 27 | 0.5922 | 0.982 | 3068 | 0.04810575 | 0.972 |
| 36302 | Homo sapiens | GSM823380 | POLR2A | None | T Lymphocyte | Blood | 39 | 0.7371 | 0.8 | 10180 | 0.2364385 | 0.9728 |
| 38342 | Homo sapiens | GSM1022949 | POLR2A | None | T Lymphocyte | Blood | 39 | 0.736 | 0.968 | 5581 | 0.08956125 | 0.973 |
| 38361 | Homo sapiens | GSM1022946 | POLR2A | None | T Lymphocyte | Blood | 38 | 0.7341 | 0.964 | 6984 | 0.1400275 | 0.975 |
| 38431 | Homo sapiens | GSM1022950 | POLR2A | None | T Lymphocyte | Blood | 39 | 0.7477 | 0.97 | 6222 | 0.09659675 | 0.971 |
| 38433 | Homo sapiens | GSM1022948 | POLR2A | None | T Lymphocyte | Blood | 39 | 0.7713 | 0.974 | 1407 | 0.03362775 | 0.9644 |
| 38454 | Homo sapiens | GSM1022945 | POLR2A | None | T Lymphocyte | Blood | 39 | 0.7491 | 0.957 | 5196 | 0.079669 | 0.9732 |
| 47435 | Homo sapiens | GSM1201946 | REST | None | T Lymphocyte | Blood | 37 | 0.6073 | 0.886 | 1840 | 0.0628475 | 0.916 |
| 44097 | Homo sapiens | GSM1056935 | RUNX1 | None | T Lymphocyte | Blood | 39 | 0.7136 | 0.886 | 1187 | 0.0181065 | 0.9504 |
| 53629 | Homo sapiens | GSM1577746 | STAT5B | None | T Lymphocyte | Blood | 37 | 0.7536 | 0.767 | 8377 | 0.1459865 | 0.95 |
| 53630 | Homo sapiens | GSM1577747 | STAT5B | None | T Lymphocyte | Blood | 37 | 0.7297 | 0.708 | 8462 | 0.1848495 | 0.9604 |
| 53631 | Homo sapiens | GSM1577748 | STAT5B | None | T Lymphocyte | Blood | 38 | 0.6881 | 0.676 | 2727 | 0.07564975 | 0.804 |
| 5195 | Homo sapiens | GSM630810 | YY1 | None | T Lymphocyte | Blood | 20 | 0.5189 | 0.783 | 3788 | 0.119051698 | 0.9708 |
